# In vitro reconstitution demonstrates the amyloid-beta mediated myelin membrane deformation

**DOI:** 10.1101/2021.08.13.456302

**Authors:** Anuj Tiwari, Sweta Pradhan, Achinta Sannigrahi, Suman Jha, Krishnananda Chattopadhyay, Mithun Biswas, Mohammed Saleem

## Abstract

Amyloid-beta (Aβ) aggregation mediated neuronal membrane deformation, although poorly understood, is implicated in Alzheimer’s Disease (AD). Particularly, whether Aβ aggregation can induce neuronal demyelination remains unknown. Here we show that Aβ-40 binds and induces extensive tubulation in the myelin membrane in vitro. The binding of Aβ-40 depends predominantly on the lipid packing defect densities and electrostatic interactions and results in rigidification of the myelin membrane in the early time scales. Furthermore, elongation of Aβ-40 into higher oligomeric and fibrillar species leads to eventual fluidization of the myelin membrane followed by extensive membrane tubulation observed in the late phase. Taken together, our results capture mechanistic insights into snapshots of temporal dynamics of Aβ-40 - myelin membrane interaction and demonstrate how short timescale, local phenomena of binding, and fibril mediated load generation manifests into long timescale, global phenomena of myelin tubulation and demonstrates the ability of Aβ-40 to demyelinate.

## INTRODUCTION

Aggregation and accumulation of Amyloid-β (Aβ)peptides in the extracellular space of neurons, forming senile plaques, is considered a hallmark of Alzheimer’s Disease (AD)(Selkoe & Hardy, 2016; Winblad *et al*, 2016). The cleavage of a transmembrane amyloid precursor protein results in the generation of two pathologically important amyloid-ß (Aβ)peptides - Aβ-40 and Aβ-42(Haass & Selkoe, 1998). While the non-aggregated forms of Aβ are known to be physiological constituents of cerebrospinal fluid, the neuritic plaques are rich in fibrillar forms of Aβ (Selkoe, 2000; Seubert *et al*, 1992). The understanding of the pathology of AD has undergone a paradigm shift over time. It is now believed that the soluble oligomeric Aβ forms are critical factors in driving the onset and progression of neuronal injury and have more potent neurotoxic effects than mature aggregates (Bucciantini *et al*, 2002; Kirkitadze *et al*, 2002). While the synaptic membrane disruption has attracted significant focus in AD research, however, the myelin membrane’s potential role remains unknown. Emerging clinical evidence strongly suggests that demyelination is an essential pathological feature that may be one of the earliest characteristics during AD progression (Dean *et al*, 2017). Despite the evidence that the soluble forms of Aβ are elevated in the white matter independent of the amyloid burden in the cortical plaque, whether Aβ aggregation can alter the myelin membrane remains elusive(Collins-Praino *et al*, 2014).

The etiology of AD remains poorly understood due to the interplay of highly complex mechanisms that govern Aβ aggregation and dynamic cellular interface. The complexity of the problem arises primarily due to - the existence of soluble aggregates that are highly heterogeneous in size, shape, and structure(Fusco *et al*, 2017); aggregation induced by seeding(Jucker & Walker, 2013); and the interplay between highly diverse physicochemical properties of lipid membranes and Aβ aggregation kinetics(Di Paolo & Kim, 2011). The fibrillogenic properties of Aβ-40 and membrane damage have been observed to be significantly correlated (Yip & McLaurin, 2001). The formation of heterogeneous ion channels observed before the plaque formation has been considered the early steps during neuronal damage formed by soluble, small-sized aggregates of Aβ-40 (Kourie, 2001). However, the inertness of Aβ-40 mature fibrils is not fully validated as both fibrils and oligomeric Aβ-40 assemblies induced a decrease in mitochondrial membrane potential in neurons (Eckert *et al*, 2008). More importantly, the same was not observed for neurons incubated with Aβ-40 monomeric forms(Eckert *et al*., 2008). Gangliosides and cholesterol have also been observed to enhance the Aβ-40 membrane interaction and influence the peptide’s penetration into the membrane(Arispe *et al*, 1993; Ji *et al*, 2002; Kakio *et al*, 2001; Sciacca *et al*, 2012b). Therefore, despite the body of reported work supporting that the amyloid cytotoxicity is majorly due to small oligomeric Aβ-40, toxic peptide’s real nature is still a matter of intense debate(Sciacca *et al*, 2018). Although the two-step mechanism involving both the pore formation and fibril mediated membrane deformation is a relatively newly proposed general mechanism behind neuronal membrane damage, however, a full understanding of the interplay of the membrane parameters, early stages of membrane-associated aggregation, and membrane deformation remains elusive(Lauwers *et al*, 2016; Sciacca *et al*., 2018; Shrivastava *et al*, 2017). More recently, studies indicate that the templated protein misfolding (seeding) could be a crucial mechanism in the initiation and propagation of Aβ as the slow primary nucleation cannot by itself account for the steep aggregation kinetics that follows(Jucker & Walker, 2013; Kane *et al*, 2000). Therefore, secondary nucleation mechanisms such as seeding result in the acceleration of amyloid fibril formation by reducing or inhibiting the lag phase of fibril formation and are now considered the major driving force in the progression of Aβ aggregation(Cohen *et al*, 2013). The current understanding is obtained mostly using small liposomes or supported bilayers composed of single/binary lipid membrane mixtures and at peptide concentrations towards the higher side in micromolar concentrations (i.e., 10-40 μM). Further, the studies reported so far did not take into account the complex lipid compositions such as myelin, the existence of phase-separation, and the secondary nucleation mechanism into account that are crucial for the initiation and progression of neuronal membrane damage. Also, there is a lack of temporal mapping of the membrane deformation induced by Aβ during its aggregation.

Here we reconstitute *in vitro* the binding of seeded Aβ on the myelin model membrane and visualize snapshots of the changes at membrane interface over a period of early, mid and late phases of Aβ aggregation spanning 24 hours, using a combination of photonic and electron microscopy, fluorescence spectroscopy, and membrane monolayer experiments. Enhanced fluidization of the myelin membrane during early binding and elongation of Aβ was found to precede the extensive membrane tubulation observed in the late phase involving fibril mediated weak phase separation. Dissection of the early binding of Aβ to myelin lipid components varying in their shape and charge revealed particularly high lipid specificity for DOPG, PI, BSM, and PIP_2_ membranes. Coarse grain MD simulations showed the highest density of lipid packing defects in the myelin membrane compared to other membranes containing diverse lipid shapes. Indeed, the binding and diffusion of Aβ and the reduction in lipid diffusion were strongest in the case of myelin membrane during the early to mid and phase, suggesting a correlation between Aβ binding and lipid packing defects at the membrane interface. Furthermore, the seeded Aβ was found to significantly deform the myelin monolayer membrane equilibrated at a bilayer equivalent pressure within the early time regime. Our findings suggest that under the seeded environment, the early binding of Aβ on the myelin membrane strongly depends on the density of the surface lipid packing defects that help drive the fibril load generation and deformation of the myelin membrane.

## RESULTS

### Binding and deformation of myelin membrane by Aβ-40

To examine whether the Aβ-40 interaction could trigger the deformation of the myelin membrane, we sought to use the reconstitution methodology to allow us to mimic myelin membrane exposed to a bulk concentration of soluble Aβ-40 and map any morphological changes in the membrane over different phases of Aβ-40 aggregation. This is particularly important as more and more emerging clinical evidence suggests that demyelination takes place before neuronal damage and the fact that elevated levels of soluble Aβ-40 are present in the white matter (Collins-Praino *et al*., 2014; Ferreira *et al*, 2020; Tse *et al*, 2018). To this end, giant unilamellar vesicles (GUVs) were reconstituted using the lipid components of the myelin membrane to mimic the outermost bilayer of the myelin sheath membrane(Calderón & DeVries, 1997) (Fig. 1). Further, to mimic the physiologically more relevant secondary nucleation mechanisms that have been suggested as a major driving force in progressive protein aggregation(Sowade & Jahn, 2017), we pre-seeded Aβ for a shorter time with oligomeric Aβ and then added to membrane and monitored over 1, 4, 12, 24 hours. To quantify different populations of soluble forms of Aβ-40 by fluorescence correlation spectroscopy, we measured the hydrodynamic radii extracted through the diffusion coefficients determined by fluorescence correlation spectroscopy, wherein the monomer to oligomer ratio of 9:1 was observed (Fig. S1, Table S1). From here on, throughout the manuscript, the early phase refers to the 0-4 hour time point, mid-phase refers to the 4 to 12 hour time point, and the late phase refers to the 12 to 24 hour time point for ease of understanding. We observed strong early binding of the Aβ-40 to myelin model membrane composed of DOPC/BSM/DOPE/PI/DOPS/Cholesterol(Calderón & DeVries, 1997). The chosen composition mimics the complexity of a myelin membrane, both in terms of the topological aspects of the lipids (i.e., a mixture of cylindrical, conical, inverted conical lipids with varying hydrophobic volumes) and surface charge (Fig. 1, Fig. S2). Aβ-40 showed significant binding to the myelin membrane starting early phase (i.e., visualized at 1 hour) as evident from the binding intensity, followed by a decrease in the binding at 4-hour time point. The early phase likely involves the binding of the population of Aβ-40 that is predominantly monomeric to lower oligomeric. However, the binding of short fibrils cannot be ruled out as it is technically challenging to quantify the same in the bound state given the highly dynamic state of aggregation. Interestingly, at the 12-hour time point again, an increase in the binding intensity was observed (Fig. 1). The variation in the binding intensity of Aβ-40 at the observed time points is likely due to the destabilization of the membrane by Aβ-40 as a result of its early phase of aggregation and subsequent opposing equilibrium forces operating within the membrane. Interestingly, at 24 hours, Aβ-40 was found to induce striking membrane tubulation at the interface, as evident from the tubulation profile of the large distorted membrane regions (Fig. 1). The quantification of the measure of tubulation observed estimated by ImageJ is detailed in the methodology section. From the observed tubular morphology, it is difficult to conclude whether these are aggregates of extracted lipid and Aβ-40 or coating of the membrane tubules by elongating Aβ-40. The microscopic observations of the myelin membrane deformation induced by Aβ-40 point to a clear indication of its aggregation mediated deformation. However, it does not convey the high-resolution structural details of the membrane-bound Aβ-40. Negative staining electron microscopy of myelin membrane mimic incubated with Aβ-40 over a time period of 4, 12, and 24 hours confirmed the fibrillar network as well as the progression of Aβ-40 induced disruption of the myelin membrane (Fig. 1). The fluorescence microscopy observation suggests that while the process of destabilization of the myelin membrane is triggered by the early binding of Aβ-40 aggregation, however, significant tubular disruption takes place only as the Aβ-40 progresses through the late fibrillar phase of aggregation. Moreover, Aβ-40 aggregation is known to be enhanced by curvature, and therefore, it might be reasonable to believe that these are indeed membrane tubules coated by Aβ-40(Terakawa *et al*, 2018).

**Fig. 1.**
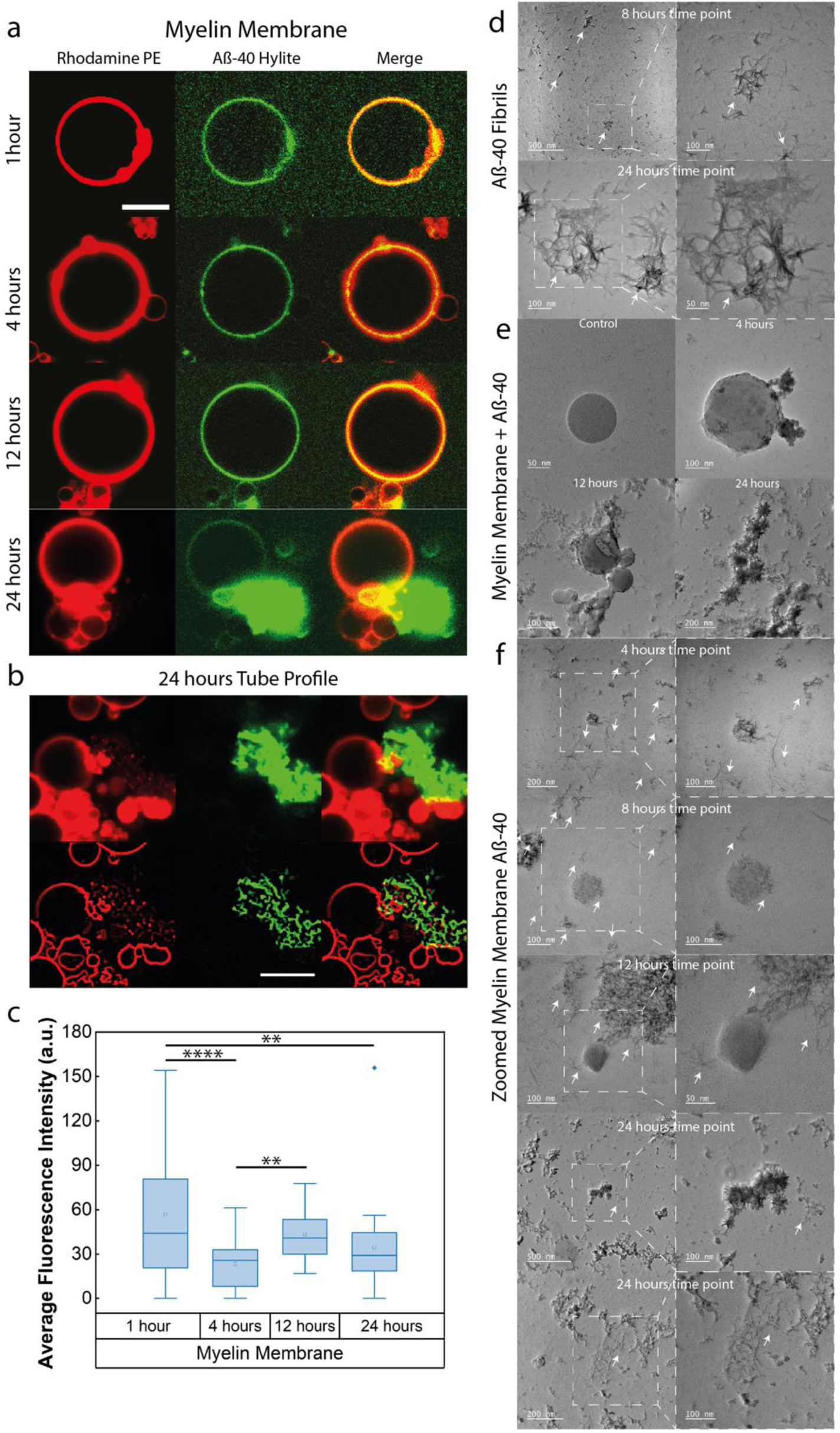
Binding and deformation of myelin membrane by Aβ-40. **a)** Temporal changes in the morphology of the GUV equatorial contour observed in confocal imaging observed at 1, 4, 12, and 24 hours with representative images shown from a pool of GUVs at each time point. GUVs are labeled with Rhodamine PE (*red channel*) and incubated with Aβ-40 doped with Hylite-488 Aβ-40 (*green channel*) under evaporation free conditions to monitor the changes in the membrane during Aβ-40 aggregation phase. **b)** Tubeness profile of the membrane morphology induced by Aβ-40 at late aggregation phase. **c)** Average fluorescence intensity of the Aβ-40 binding at the equatorial plane of GUV observed at 1, 4, 12, and 24-hours time point. The number of GUVs screened in the box plot is n= 35 from three independent experiments. The symbols **, **** indicate p values of ≤ 0.01, 0.0001, respectively, calculated by one-way ANOVA followed by Bonferroni’s multiple comparison test. **d)** TEM micrographs of Aβ-40 aggregation at 8 and 24 hours with marked inset for the zoomed images on the right. **e)** TEM micrographs of temporal changes in the membrane morphology induced by Aβ-40 aggregation imaged over 24 hours. **f)** TEM micrographs of temporal changes in the membrane morphology induced by Aβ-40 aggregation imaged over 24 hours with marked inset for the zoomed images on the right. White arrows in the TEM micrographs mark the fibrils. The scale bar for confocal microscopy images is 10 μm.

### The interplay of lipid specificity and fibrillation of Aβ-40

Considering that several lipid components constitute the myelin membrane, we next asked whether there is specificity for diverse lipid geometry and the head group that could play an important role in dictating Aβ-40 binding and aggregation. The shape of lipids depends on the aspect ratio of their headgroups and acyl chains that determine their packing, the degree of local defects, and spontaneous curvature within the membrane(McMahon & Boucrot, 2015). To evaluate this, we set out to investigate microscopic visualization of the temporal changes in membrane upon binding by Aβ-40. To this end, we first focused on the early phase of its interaction and aggregation corresponding to a time scale of the first hour. Interestingly, while no significantly visible early binding of Aβ-40 was observed in DOPC (conical/zwitterionic lipid) membrane in the first hour(Strandberg *et al*, 2012) (Fig. 1a), homogenous binding was observed in the case of DOPG (cylindrical/negatively charged lipid) and PI membrane (inverted conical/negatively charged lipids (Fig. 2a, Fig. S3).

**Fig. 2.**
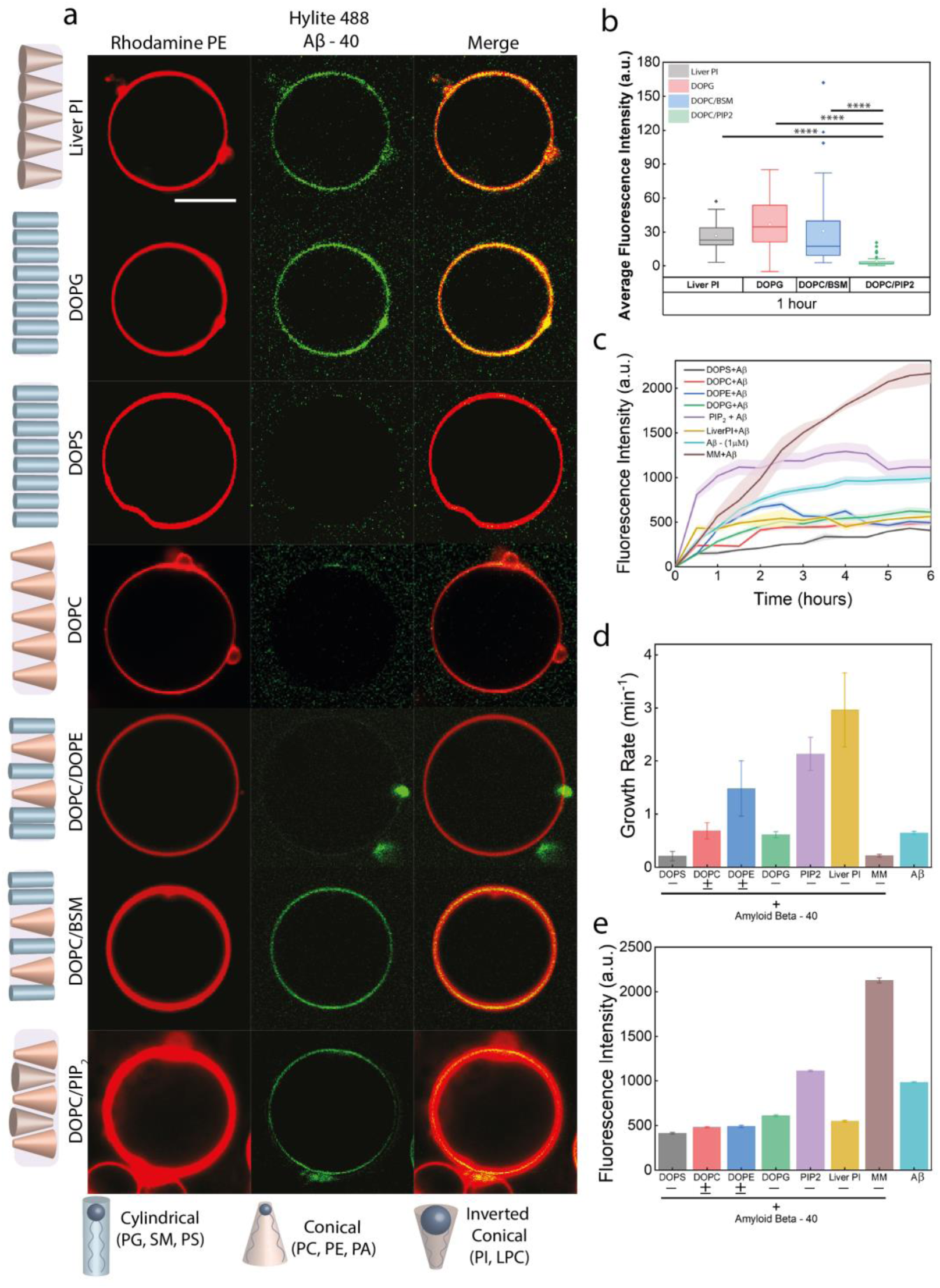
Lipid specificity of the Aβ-40 early binding and fibrillation. **a)** GUVs labeled with Rhodamine PE (red channel) were incubated with Aβ-40 doped with Hylite-488 Aβ-40 (green channel) to monitor binding of the peptide within 1 hour (early phase). Early binding of Aβ-40 was observed in the case of PI, DOPG, DOPC/BSM, and DOPC/PIP2 membrane. No binding of Aβ-40 seen on DOPS and DOPC/DOPE membranes. **b)** Average fluorescence binding intensity of Aβ-40 on membranes with different head groups. The number of GUVs screened for each condition in the box plots is n= 35 from three independent experiments. The symbol **** indicates p values of ≤ 0.0001, calculated by one-way ANOVA followed by Bonferroni’s multiple comparison test. **c)** ThT fluorescence assay to monitor the effect of lipid head groups on the aggregation kinetics of Aβ-40. **d)** Effect of lipid head groups on the growth rate of Aβ-40 aggregation quantified within in the early phase of aggregation time scales i.e., 30 minutes). **e)** Effect of lipid head groups on the sustained aggregation of Aβ-40 during the late phase (saturation phase) of aggregation kinetics, i.e., 6 hours). +/- refer to the net charge on the membrane (i.e, negative or zwitterionic). The scale bar for confocal microscopy images is 10 μm.

Further, no binding was observed in the DOPC/DOPE (3:2) membrane composed of both cylindrical and conical lipids with an overall neutral charge(Bera *et al*, 2020) (Fig. 2a). Surprisingly, Aβ-40 was found to bind the DOPC/BSM (3:2) membrane composed of only cylindrical lipids but with different acyl chain length (hydrophobic mismatch) and neutrally charged, more intensely, compared to the DOPC/PIP2 (3:2) membrane composed of both cylindrical and inverted conical lipids with a net negative charge (Fig. 2a). We further examined the influence of the constituent lipid components (DOPC, DOPE, DOPS, PIP2, PI, DOPG, and myelin membrane) on aggregation kinetics of Aβ-40 by incubating 1μM Aβ-40 with membranes composed of 200μM of different lipids. Examining the growth rate measured for the initial surge in the fluorescence intensity observed within a time window of 30 minutes, amongst all the lipid membranes, PI triggered the fastest fibril formation followed by PIP2, both of which are inverted conical in shape (Fig. 2c,d). Both zwitterionic cylindrical DOPC and negatively charged cylindrical DOPG were found to have no significant effect on the fibrillation in the early phase kinetics as evident from extracted growth rates (Fig. 2d). However, looking at the overall aggregation kinetics, at late or saturated phase, DOPC tends to slow down the Aβ-40 aggregation. This observation is consistent with previous reports wherein the DOPC membrane was found to slow down the aggregation kinetics of Aβ-40(Hellstrand *et al*, 2010; Sahoo *et al*, 2019) (Fig. 2c, 2e).

Interestingly, despite the early binding of Aβ-40 on the myelin membrane, the observed growth rate was lowest amongst all, hinting at slow fibrillation (Fig. 2d). The myelin membrane was also found to facilitate Aβ-40 fibrillation most strongly and sustain it over a longer duration, as evident from the ThT fluorescence intensity observed at the plateau (Fig. 2e). Amongst all the membranes, only PIP2 was found to facilitate fibrillation as evident from higher plateau (Fig. 2e). PIP2 membrane with a net negative charge, the apparent facilitation of aggregation of Aβ-40 which itself is a negatively charged peptide seems counter intuitive. This affinity of the peptide for the negatively charged bilayer could be attributed to the coulombic interaction of positive residues present in Aβ-40(Kang & Sun, 2020; Yang *et al*, 2021). The differential binding of Ab-40 to single and binary lipid compositions such as between DOPC & DOPS (No early binding of Aβ) and DOPC/BSM, DOPC/PIP2 (early binding of Aβ was observed) were intriguing and led us to conclude there are more parameters involved besides the electrostatic interactions that might be important for the binding. And one such parameter that was strikingly different for different components of the myelin mimic was the lipid shape. Thus, we set out with a hypothesis that given the complexity of myelin membrane and the diverse shapes of its lipid components, can lipid defects play any role besides the electrostatic interactions in myelin membrane deformation?

### Myelin membrane contains higher lipid packing density defects

We hypothesized that the interplay of lipid packing defects and electrostatics drive the early binding, fibrillation of Aβ-40 and subsequent deformation of the membrane. To quantify the lipid packing defect density, we used coarse-grained molecular dynamics simulation. The following four membrane surfaces containing lipid compositions with different shapes, hydrophobic volumes, and charge were adopted to mimic different degrees of lipid packing defects - i) zwitterionic conical lipids (DOPC), ii) zwitterionic conical and cylindrical lipids (DOPC/BSM/Chol), iii) negatively charged inverted conical, zwitterionic conical and cylindrical lipids (DOPC/BSM/PIP2/Chol), iv) a more complex surface containing negatively charged inverted conical, zwitterionic conical and cylindrical lipids (myelin). We used PackMem(Gautier *et al*, 2018) to quantify the lipid packing defects that follow the Cartesian grid system for mapping the membrane surface where the grid dimension is set to 1 Å x 1 Å. This tool computes the defects by characterizing them into deep and shallow defects. The deep defects represent the voids created due to the presence of aliphatic atoms deeper than d Å (where the value of d is set to 1Å) below the central atom of glycerol, whereas shallow defects represent the accessible aliphatic atoms that are less than d Å below the central atom of glycerol and all types represent the combination of both the deep as well as shallow defects(Gautier *et al*., 2018). Using PackMem execution upon each of the membrane systems, we produced a plot comparing the value of defect constant (π) for the three types of defects (deep, shallow, and all) shown in the figure below (please see methods for details of CGMDS):

The higher the π constant, the more abundant and larger the packing defects(Gautier *et al*., 2018). Overall, of the four bilayer systems studied, the myelin membrane has more numbers of packing defects, and out of deep and shallow defects, shallow defects are more abundant. The scale of the observed variation in the lipid packing defect densities of chosen membrane surfaces is in the range of 3-5 Å holds significant relevance in the biological regime.

### The interplay of lipid packing defects and electrostatics drives Aβ-40 fibrillation and membrane deformation

To validate our hypothesis experimentally, we next questioned if the observed differences in the lipid packing defect densities might be the driving forces for the degree of early binding and subsequent deformation by Aβ-40. Since Aβ-40 aggregation is a slow process, we monitored the fate of Aβ-40 binding and changes induced in membrane morphology on a time scale of 24 hours and captured the snapshots at early (1-4 hour), mid (4-12 hour), and late aggregation phases (12-24 hour) (i.e., at 1 hour, 4 hours, 12 hours, and 24 hours respectively) (Fig. 4, Fig S3-5). We observed that in the case of DOPC, Aβ-40 was weakly membrane-bound at 4 hours with no microscopically visible deformation of the membrane, as evident from the contour on the equatorial plane of the GUV (Fig. 4). Striking deformation of the membrane into large tubular structures was observed at 24 hours, followed by visualization of a pool of GUVs that either has intense membrane tubulation or completely ruptured membrane in the late phase (around 24 hours) (Fig. 4a-b).

We next investigated the binding of Aβ-40 to the membrane composed of DOPC, BSM, and Cholesterol (4:4:2), accommodating a relatively lesser amount of lipid packing defects compared to the DOPC membrane. Interestingly, unlike the DOPC/BSM membrane, which showed early binding of Aβ-40 (Fig. 1a), negligible to weak binding of Aβ-40 to DOPC/BSM/Chol membrane was observed in the early phase (1-4 hours) followed by increased binding and mild tubulation during the mid-phase (4-12 hours) (Fig. 4c-d). Similar to the DOPC membrane, a pool of tubulating and collapsed membrane was observed at late-phase (24 hours) (Fig. 4c-d), although the tubulation was not as strong as in the case of DOPC membrane (Fig. 4a). We reasoned that the presence of cholesterol is expected to increase the lipid packing density and stiffen the membrane containing unsaturated lipids, thereby reducing the membrane tubulation(Chakraborty *et al*, 2020). We then wondered about the consequence of adding a negatively charged inverted conical lipid component to the DOPC/BSM/Chol membrane, and therefore, we added 10 mol% PIP2 to the above mix while decreasing DOPC, keeping the BSM unchanged, and increasing the cholesterol levels ((DOPC/BSM/Chol/PIP2 (2:4:3:1)). Interestingly, strong binding of Aβ-40 was observed in the presence of PIP2 during early to mid-phase, even though relatively excess cholesterol was present in the membrane. Likewise, strong tubulation of the membrane was observed during the late phase (Fig. 4e-f). No binding of Aβ-40 was observed from the early to the late phase when PIP2 in the DOPC/BSM/Chol/PIP2 membrane was replaced with DOPG (a negatively charged cylindrical lipid) that results in a reduction in lipid packing defects (Fig. 4g). This has also been found in related membrane composition where DOPG likely interacts with neighboring lipid molecules by hydrogen bonding. This hydrogen bonding is between glycerol moiety of DOPG and the phosphate oxygen of the neighboring phospholipid which leads to ordering of the membrane(Greiner *et al*, 2009). The early membrane binding of Aβ-40 could be predominantly driven by the lipid packing defects and the limiting bulk peptide concentration; however, the polar interactions should be essential for the stabilization of the interactions. This is evident from the lack of early binding of Aβ-40 to DOPC that has significant lipid packing defects but not sufficiently strong electrostatic forces (Fig. 3, 4a). Further, considering the membrane deformation and comparing the membrane binding induced by Aβ-40 over time across the membrane conditions investigated, the degree of binding of Aβ-40 on membranes decreases in the following order - myelin membrane (cylindrical lipids with height mismatch, inverted conical and conical lipids) > DOPC/BSM/Chol/PIP2 (conical, cylindrical, inverted conical lipids) > DOPC/BSM/Chol (conical and cylindrical lipids) > DOPC (conical lipids)(Box plot, Fig. S6-8). Together, the above observations suggest that although the early binding and fibrillation of Aβ-40 depends on the interplay of both lipid geometry (that defines local lipid packing defects) and electrostatics. However, lipid packing defects could be the predominant factor amongst the two that dictates both the kinetics of Aβ-40 binding and membrane deformation.

**Fig. 3.**
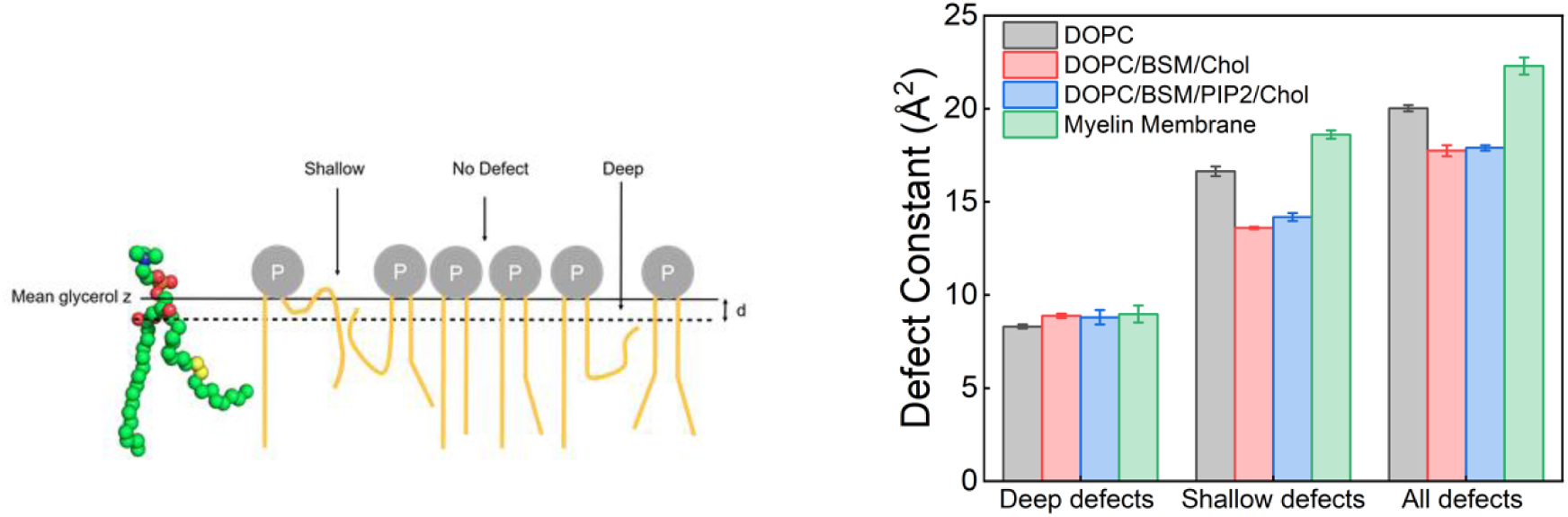
Simulation-based quantification of the lipid packing defect constants. **a)** An illustration to provide a lateral view of shallow, deep, and no defects at the membrane surface. **b)** Histogram of the defect constant for deep, shallow, and all (both deep and shallow defects) defects for each of the four conditions, namely, DOPC (red), DOPC/BSM/Chol (green), DOPC/BSM/Chol/PIP_2_ (blue), Myelin membrane (cyan).

**Fig. 4.**
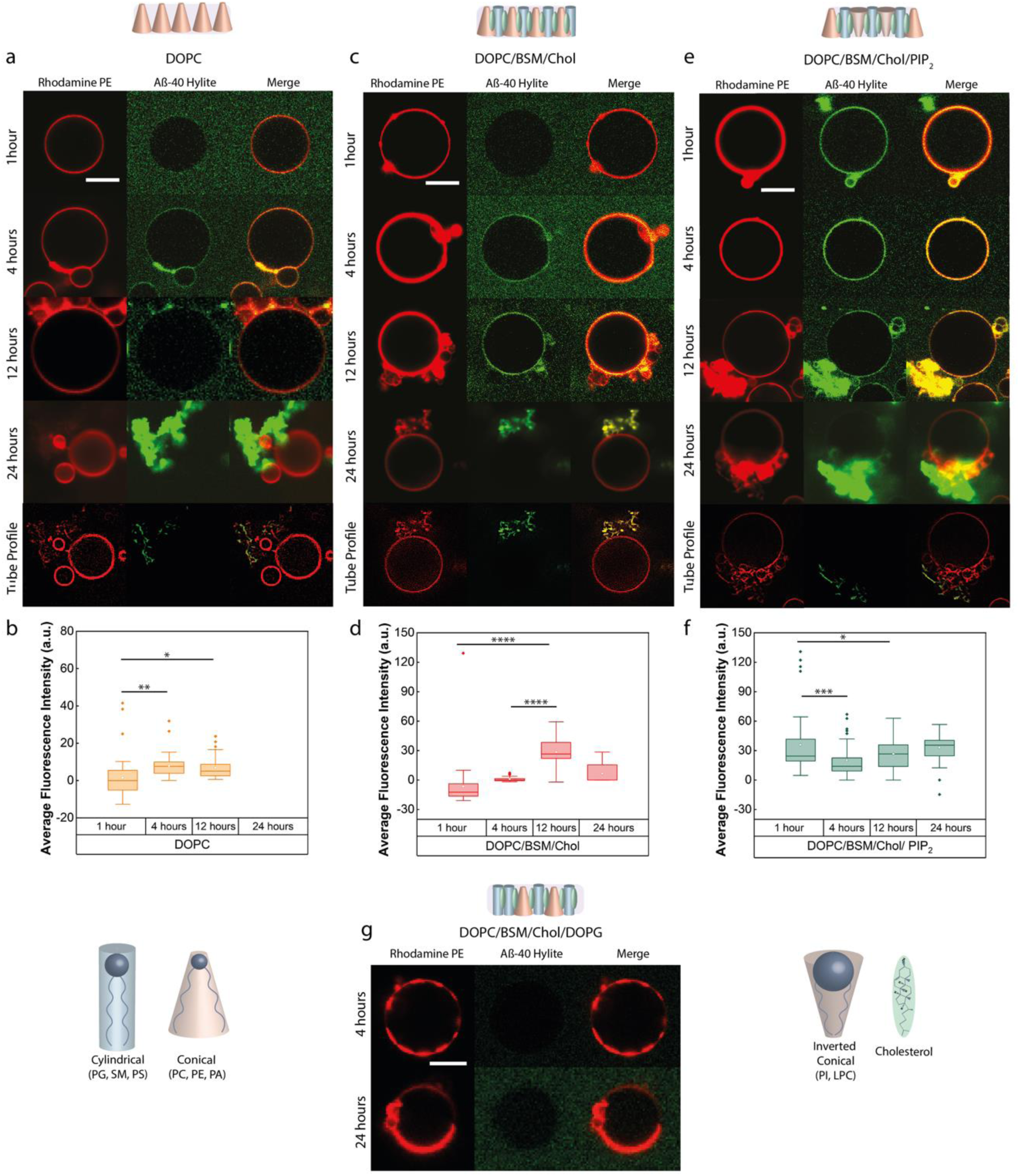
Effect of lipid packing defects and electrostatics on Aβ-40 binding and membrane deformation. **a)** GUVs of DOPC lipid doped with 1% Rhodamine PE (Red channel) incubated with Aβ-40 doped with Hylite-488 Aβ-40 (Green channel) and monitored temporally at 1, 4, 12 and 24 hours for changes in binding intensity. Tube profile was extracted at the 24-hour time point to visualize the deformations induced by Aβ-40 over time. **b)** Box plot showing weak to no binding of Aβ-40 to the DOPC GUV population observed at an early time point (1 hour), whereas, at later time points the binding increased and reached a plateau. **c)** Temporal monitoring of GUVs of DOPC/BSM/Chol lipid in the ratio 4:4:2 doped with 1% Rhodamine PE (Red channel) and incubated with Aβ-40 doped with Hylite-488 Aβ-40 (Green channel) at 1, 4, 12 and 24 hours for changes in binding intensity. The tube profile was extracted at the 24 hour time point to visualize the deformations induced by Aβ-40 over time. **d)** No binding was seen at the early time point (1 hour) for the DOPC/BSM/Chol (4:4:2) GUV population, with a steady rise in intensity at later time points. **e)** GUVs of DOPC/BSM/Chol/PIP_2_ lipids in the ratio 2:4:3:1 doped with 1% Rhodamine PE (Red channel) incubated with Aβ-40 doped with Hylite-488 Aβ-40 (Green channel) and monitored temporally at 1, 4, 12 and 24 hours for changes in binding intensity. The tube profile was extracted at the 24-hour time point to visualize the deformations induced by Aβ-40 over time. **f)** Box Plot of the binding intensity of Aβ-40 to the GUV population of DOPC/BSM/Chol/PIP_2_ (2:4:3:1) showing a strong early binding sustained till longer time points. **g)** Temporal monitoring of DOPC/BSM/Chol/DOPG (2:4:3:1) GUVs doped with 1% Rhodamine PE (Red channel) incubated with Aβ-40 doped with Hylite-488 Aβ-40 (Green channel) at 4 and 24 hours showed no binding at early and late time points. The number of GUVs screened at each time point for each condition in the box plots is n= 35 from three independent experiments. The symbols *, **, ***, **** indicate p values of ≤ 0.05, 0.01, 0.001, 0.0001, respectively, calculated by one-way ANOVA followed by Bonferroni’s multiple comparison test. The scale bar for confocal microscopy images is 10 μm.

### Contribution of lipid shape in the absence of charge for Aβ-40 binding

We then looked into the role of coupling between proportions of different lipid shapes (i.e., a ratio of cones to cylinder) and cholesterol, in the absence of negative charge, that might affect the binding of Aβ-40. To test this, we modulated the ratios of DOPC (conical lipid), BSM (cylindrical lipid), and cholesterol (Fig. S9). We observed significant early binding of Aβ-40 on DOPC/BSM membranes in the absence of cholesterol both at a ratio 5:5 as well as 3:2 (Fig. S9 and Fig. 2). Interestingly the binding efficiency of Aβ-40 was found to vary as the proportions of conical/cylindrical lipid changed, suggesting that not only the lipid shape is important but also the proportions of different lipid shapes might influence the lipid packing defects and subsequent binding of Aβ-40. Evidently, in the presence of an equal proportion of cholesterol, the reversal of the ratios of DOPC: BSM from 2:6 to 6:2 results in a significant weakening of the Aβ-40 early binding on the membrane (Fig. S9). Interestingly, even for DOPC/BSM/Cholesterol ratio of 3:3:4 with double the proportion of cholesterol, early binding of Aβ-40 is observed to the liquid disordered regions of the phase-separated membrane (Fig. S9). This further reinforces our hypothesis that lipid geometry that dictates packing defects contribute predominantly in the initial binding of Aβ-40.

### Aβ-40 drives myelin deformation through its fluidization

We reasoned that the observed differences in early and late binding of Aβ-40 should also reflect in the fluidity changes in the membrane at a comparable time point. Irrespective of the density of lipid packing defects, Aβ-40 seems to bind to most membrane conditions by the 4 hour time points (early to mid-phase) (Fig. 4). Thus, we next probed the fluidity changes in the membrane around 4 hours, owing to the aggregation of Aβ-40. Steady-state fluorescence anisotropy measurements report the global changes in the fluidity of the lipid membrane. The changes in membrane fluidity were quantified by reconstituting a potentiometric styryl membrane probe di-8-ANEPPS within membrane vesicles, whose fluorescence anisotropy responds to the changes in the dynamics of the membrane environment(Le Goff *et al*, 2007; Starke-Peterkovic *et al*, 2006). We also found that the trend remained largely unchanged till 8 hours (Fig S11). The measure of anisotropy obtained for different membrane conditions in the presence of Aβ-40 was normalized by the anisotropy obtained for respective control membrane for each condition, such that all the control values were scaled to 100 for ease of comparison (marked as a red line across the graph, Fig. 5).

**Fig. 5.**
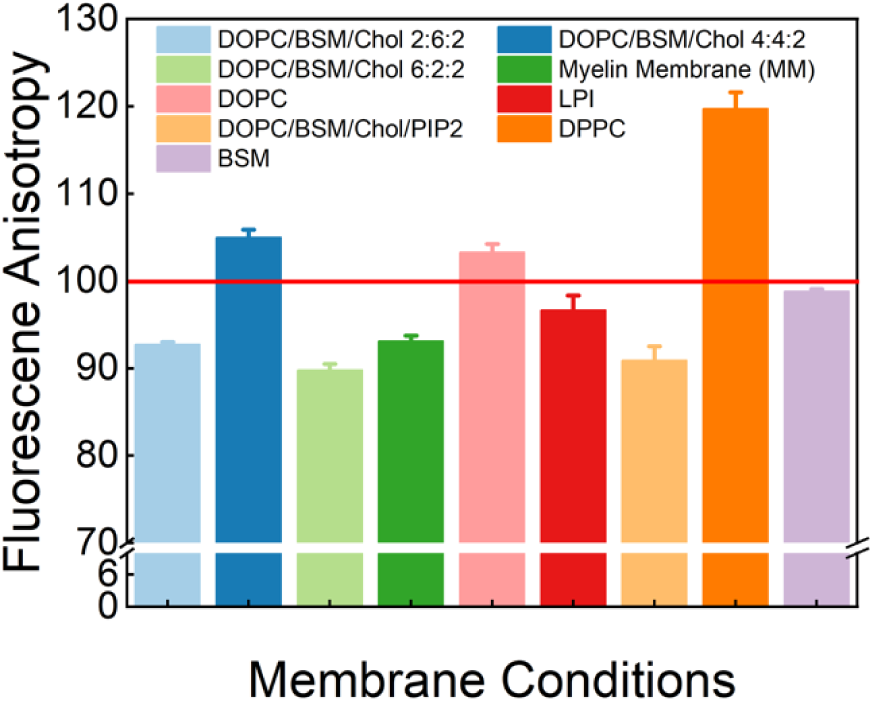
Changes in membrane fluidity by Aβ-40. Steady-state anisotropy measurements were performed on different membrane conditions that include membranes with variable DOPC/BSM content keeping cholesterol constant (2:6:2, 4:4:2, 6:2:2), along with single lipid membrane compositions like DOPC, BSM, LPI, DPPC, and multicomponent membranes of myelin and DOPC/BSM/Chol/PIP_2_ incubated with Aβ-40. Controls without Aβ-40 for each membrane condition were normalized to 100, denoted by the horizontal red line traversing the plot. The data shown represent the ± SEM extracted from three independent experiments.

We observed that the binding of Aβ-40 to both myelin and DOPC/BSM/Chol/PIP2 membrane results in a reduction in the fluorescence anisotropy of the probe, indicating an increase in fluidization. On the contrary, binding to DOPC/BSM/Chol (4:4:2 mol ratio) and DOPC membranes results in an increase in the fluorescence anisotropy of the probe, suggesting a decreased fluidization (Fig. 5). Further, PI and DOPC/BSM membranes also were found to be fluidized, as evident from the decrease in the observed anisotropy. Modulating the ratios of DOPC and BSM within the DOPC/BSM/Chol membrane (2:6:2 and 6:2:2) showed only a little difference in the enhanced fluidity. DOPC membrane seems to have rigidified by the binding of Aβ-40 as suggested by the increased anisotropy. To further confirm the correlation between lipid packing density, Aβ-40 binding, and subsequent increase or decrease in fluidization, we checked the effect of Aβ-40 binding on the fluidity of DPPC (saturated membrane). We observed that a significant reduction in the fluidity of the DPPC membrane was observed. This is in support of a previous study wherein it was shown that the gel phase rigid domains of DPPC may act as a platform for Aβ enrichment that might further decrease the fluidity of the membrane(Choucair *et al*, 2007). Taken together, sustained binding and fibrillation of Aβ-40 during the mid-phase results in fluidization of the membrane as evident from fluorescence spectroscopy shown in Fig. 5. This leads us to conclude that albeit the shape and charge of the lipids in the membrane play an important role in the early binding of the peptide, the process progresses to subsequent disruption through fluidization of the membrane.

### Dynamics of lipid and Aβ-40 diffusion during Aβ-40 growth

Another indication of the nature of binding of Aβ-40 and the effect on the membrane lipid diffusion can be drawn from the fluorescence recovery after photobleaching (FRAP) curves of fluorescently labeled Aβ-40 bound to the membrane at different time points. FRAP curves would suggest if the membrane-bound Aβ-40 is in a rigid immobile or a mobile structural arrangement as well as the consequent effect on the membrane lipid diffusion. We, therefore, investigated the fluorescence recovery of Aβ-40 on select membrane conditions of different lipid topology and charge as chosen earlier, at 12 hours and 24hour. The reason for selecting mid to late phase of fibrillation (i.e., 12 and 24-hour time points) is based on the observation that the predominant pool of GUVs show comparable binding at 12 and 24 hours and therefore allow bleaching of a region of interest (ROI) on the GUV equatorial plain (Fig. 6, Table S2). We observed complete recovery of fluorescent signal from Aβ-40 for the myelin membrane (containing highest lipid packing defects) at 12 hours, suggesting a dynamic interaction of Aβ-40 at the membrane interface. Photobleaching at 24hour could only bleach 35-40% of the fluorescence at the ROI, likely, due to dense coating of Aβ-40 fibrils on the membrane, which was recovered fully (Fig. 6 a-b, d-e). Likewise, ∼ 90% of the Aβ-40 fluorescence was found to recover upon photobleaching of the membrane containing moderate defects (DOPC/BSM/Chol/PIP2) at 12 hours, which eventually decreased three-folds at 24 hours (Fig. 6 a-b, d-e). No significant fluorescence recovery of Aβ-40 was observed in the case of the membrane with the least defects (PC/SM/Chol) both at 12 and 24 hours (Fig. 6 a-b, d-e). Aβ-40 seems to be in a highly dynamic and deformative interaction with the myelin membrane resulting in significant extraction of lipid tubules. This, in turn, leads to a generation of free space on the GUVs, allowing continuous binding as evident from recovery of Aβ-40 signal at 24 hours. This also hints at a likely enhancement of fluidization of the myelin membrane. Indeed, monitoring the FRAP curves of the myelin membrane lipid suggests a ∼ 20 % increased recovery in fluorescence at 24 hours compared to that at 12 hours, after photobleaching (Fig. 6c, f). A three-fold drop in the fluorescence recovery in the lipid channel at 24 hours compared to 12 hours, in the case of DOPC/BSM/Chol/PIP2 membrane, suggests a restricted movement of the lipids and more stable interfacial interaction (Fig. 6c, f). Finally, in the case of the DOPC/BSM/Chol (membrane with least amounts of defects), no significant change in the fluorescence recovery signal in the lipid channel is observed at 12 and 24 hours (Fig. 6c, f).

**Fig. 6.**
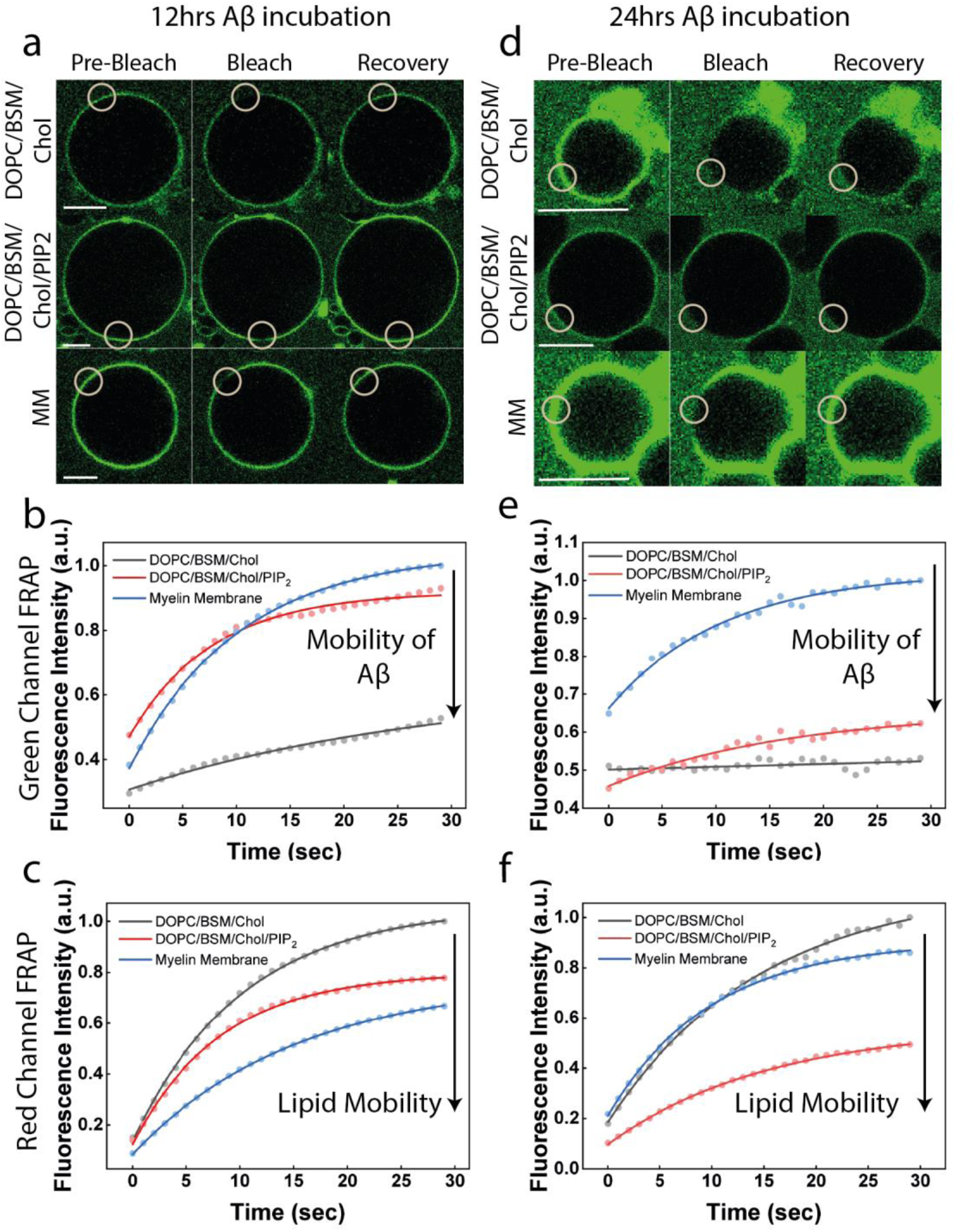
Changes in the diffusion of Aβ-40 and lipid membrane during aggregation. **a)** Representative fluorescence recovery after photobleaching (FRAP) images of the green channel (Aβ-40 doped with Hylite-488 Aβ-40) showing the Pre-bleach, bleach, and Recovery of the three conditions (DOPC/BSM/Chol (4:4:2), DOPC/BSM/Chol/PIP_2_ (2:4:3:1) and Myelin membrane) marked with a white circle to denote the region of interest in each case at the 12 hour time point. **b)** Normalized fluorescence recovery curves after photobleaching for Aβ-40 bound (green channel - peptide) to the above mentioned three membrane conditions at the 12-hour time point. **c)** Normalized fluorescence recovery curves after photobleaching for Aβ-40 bound (red channel – lipid membrane) to the above mentioned three membrane conditions at the 12-hour time point. **d)** Representative fluorescence recovery after photobleaching (FRAP) images of the green channel (Aβ-40 doped with Hylite-488 Aβ-40) showing the Pre-bleach, bleach, and Recovery of the three conditions (DOPC/BSM/Chol (4:4:2), DOPC/BSM/Chol/PIP_2_ (2:4:3:1) and Myelin membrane) marked with a white circle to denote the region of interest in each case at the 24 hour time point. **e)** Normalized fluorescence recovery curves after photobleaching for Aβ-40 bound (green channel - peptide) to the above mentioned three membrane conditions at the 24-hour time point. **f)** Normalized fluorescence recovery curves after photobleaching for Aβ-40 bound (red channel – lipid membrane) to the above mentioned three membrane conditions at the 24-hour time point. All FRAP curves in black lines are for DOPC/BSM/Chol (4:4:2), red lines are for DOPC/BSM/Chol/PIP_2_ (2:4:3:1), and blue lines represent curves for the myelin membrane. Each curve is a mean of 5 independent experiments. The scale bar for FRAP images is 10 μm.

### Aβ-40 mediated changes in phase behavior and compressibility modulus of membrane monolayer at short timescale

Bilayer experiments allowed us to probe the long time-scale phenomena (i.e, from 1 hour to 24 hours), that cannot capture the molecular aspects of interaction during earliest time scales. We therefore next aimed to investigate the short time-scale local phenomena capturing the molecular events of the earliest binding as well as changes in the mechanical properties of the membrane within 1 hour. To address this, we used two-dimensional models of a biological membrane, i.e., Langmuir monolayers that are highly sensitive tools to study mixing behavior, binding/insertion(Chi *et al*, 2008; Stefaniu *et al*, 2014; Yang *et al*, 2016). One way to identify potential specific interactions between Aβ-40 and complex membrane models is to examine the Surface Pressure-Area (*π-A*) isotherms of the membrane in the absence and presence of Aβ-40 at the air/water interface. The isotherms capture the modulation of the phase behavior and collapse pressure of the membrane induced by Aβ-40 interaction upon compression of a free-standing monolayer, particularly in the early time scales (within 30 minutes). We observed that the *π-A* isotherm of the myelin model membrane in the presence of Aβ-40 shifts towards right starting at a surface pressure of 15mN/m, indicating an increase in the area per molecule in comparison to the control myelin monolayer isotherms. This led us to conclude that Aβ-40 tends to get incorporated in the monolayer, which justifies the shift driven by the expulsion of some lipids out of the monolayer observed by the differences in the monolayer collapse pressure (*blue and light blue isotherms*, Fig. 7a-b). An interesting difference in the behavior of the DOPC/BSM/Chol/PIP2 membrane in the presence and absence of Aβ-40 (*green and light green isotherms*) is noteworthy. Although both the isotherms appear similar until the collapse pressure is approached, however, a distinct plateau is observed indicative of the coexistence of a liquid-expanded to liquid-condensed (L_e_-L_c_) region. A slight shift towards the left is observed at the coexistence plateau, indicating a decrease in the area per molecule. This observation hints at squeezing out of the lipids from the monolayer as a result of the crowding on the head groups due to the condensing effect of Aβ-40 interaction with the lipid monolayer. This is further corroborated by the difference in the collapse pressure indicative of material loss during the compression.

**Fig. 7.**
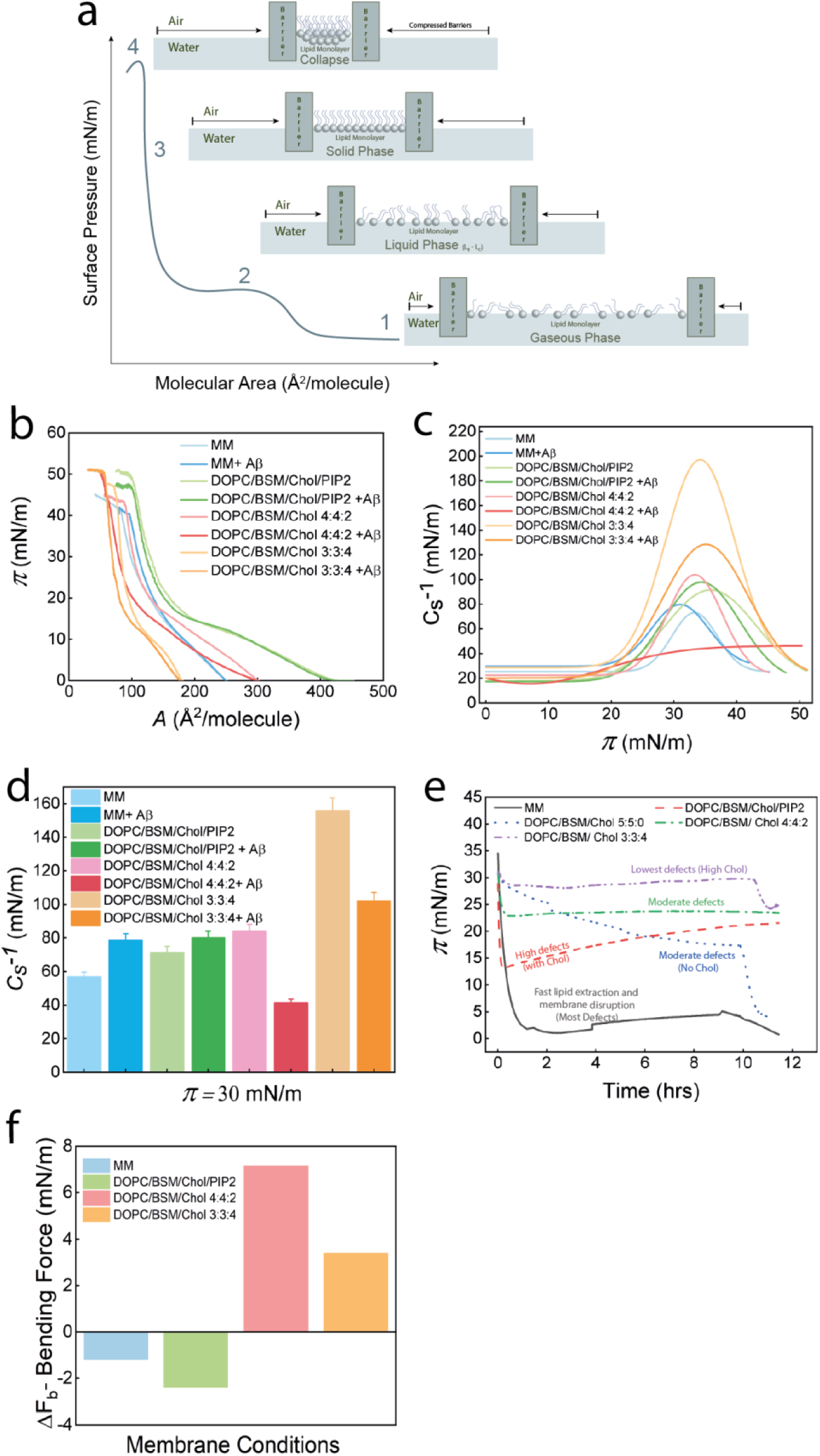
Phase behavior, compressibility modulus, and load generation induced by Aβ-40. **a)** Illustration of a typical surface pressure (*π*)– mean molecular area (*A*) isotherm with a visual representation of the molecular alignment of the lipid molecules at different phases of the isotherm. **b)** *π - A* isotherm for the different membrane conditions that include myelin membrane with and without Aβ-40 (blue and light blue), DOPC/BSM/Chol/PIP2 (2:4:3:1) membrane with and without Aβ-40 (green and light green), DOPC/BSM/Chol (4:4:2) membrane with and without Aβ-40 (pink and light pink) DOPC/BSM/Chol (3:3:4) membrane with and without Aβ-40 (orange and beige) at 25°C. **c)** Compressibility modulus (*C_s_^-1^*) *–* surface pressure (*π* ) curves for each control monolayer model as well as in the presence of Aβ-40 are shown in the graph following the same labelling order and color scheme for the membrane conditions as above. **d)** Compressibility moduli (*C_s_^-1^*) at surface pressure of 30 mN/m (at the bilayer equivalence pressure) is shown in the graph with the same labelling order and color scheme for the membrane conditions as above. **e)** Surface pressure (*π* )- Time plot for different membrane conditions that include myelin membrane (black line), DOPC/BSM/Chol/PIP2 (2:4:3:1) (red line), DOPC/BSM/Chol (5:5:0) (blue line), DOPC/BSM/Chol (4:4:2) (green line), DOPC/BSM/Chol (3:3:4) (purple line) at 25°C for 12 hours. **f)** Bending force (ΔF_b_), a measure of deformability, of different membrane conditions is shown in the histogram.

We then looked at the mixing behavior of the DOPC/BSM/Chol (4:4:2) membrane and Aβ-40. A left shift quite early in the isotherm was observed seen just at the start of the L_e_ phase indicating Aβ-40 induced condensation upon compression. The collapse of the membrane condition with Aβ-40 in the subphase happens to be higher (∼ 50 mN/m) than the control without Aβ-40. This condensing effect caused by Aβ-40 stabilized the solid phase to a greater extent indicated by an increase in the surface pressure corresponding to the collapse of the membrane models. The higher surface pressure collapse of these membranes could also be attributed to Aβ-40 having a lesser degree of affinity to these membranes, albeit having enough affinity to non-disruptively condense and stabilize the membrane. A similar observation was found for a membrane containing DOPC/BSM/Chol in 3:3:4.

The difference in the observed initial binding and phase behaviour of the membrane should manifest into changes in its mechanical properties. We therefore quantified the changes in the elastic compressibility modulus (*C_s_^-1^*) of the different membranes induced by the Aβ-40 interaction as described earlier (See methods for details). This parameter reports the in-plane elasticity (compressibility modulus) of the monolayer films and is also correlated to the different phases of the compression isotherms, namely Gaseous (G), Liquid expanded (LE), Liquid condensed (LC), and Solid (S). *C_s_^-1^* of 12mN/m to 100 mN/m corresponds to the LE region, *C_s_^-1^* ranging from 100-250 mN/m corresponds to LC, and *C_s_^-1^* of 250 mN/m and above corresponds to S(Broniatowski *et al*, 2010). Particularly, the *C_s_^-1^* of a given membrane monolayer at surface pressure (π) of 25-30 mN/m is considered to reflect the elasticity of a bilayer membrane as the lateral pressure and mechanical properties of monolayer are known to be similar to that of a bilayer at this pressure(Brockman, 1999; Brown & Brockman, 2007). The *C_s_^-1^* of the myelin membrane was close to ∼ 70 mN/m, which reaches a value in excess of ∼ 80 mN/m upon the interaction of Aβ-40. Similarly, in the case of DOPC/BSM/Chol/PIP2 membrane, the *C_s_^-1^* was found to be ∼ 85 mN/m, which slightly increased to ∼100 mN/m upon Aβ-40 interaction.

Together, the observed slight increase in the *C_s_^-1^* suggests that the interaction of Aβ-40 with the above two-membrane conditions decreases the elastic behavior of the membranes, evident from the observation that the two membrane conditions collapse within the LE phase. Interestingly, in the case of DOPC/BSM/Chol (4:4:2) membrane, the *C_s_^-1^* peaked at the value bordering ∼100 mN/m that decreased to around ∼50 mN/m upon the interaction of Aβ-40. This means that on interaction with Aβ-40, the membrane becomes more compressible. The DOPC/BSM/Chol (3:3:4) membrane was found to be the most inelastic of all the membrane composition used, reaching a *C_s_^-1^* in excess of ∼200 mN/m, that dropped to ∼130 mN/m upon the interaction of Aβ-40. It is noteworthy that both the membrane conditions showed a drastic decrease in their *C_s_^-1,^* which translates into more elastic behavior of the membranes. Membrane monolayers at a surface pressure of 30-35mN/m are considered to have a similar lateral pressure and mechanochemical properties of a lipid bilayer. Thus, analyzing the trends of the *C_s_^-1^* values for each condition at 30mN/m gives a biologically closer understanding of the changes in the elastic behavior of the membrane induced by interacting Aβ-40 that showed the same trend as observed for the peak values of *C_s_^-1^* (Fig. 7c-d). The base compressibility modulus of the myelin membrane was∼ 56 mN/m which increases to ∼78mN/m in the presence of Aβ-40, which translates to around approximately 38% increase in compressibility modulus of the model or, in other words, decrease in the monolayer compressibility. The DOPC/BSM/Chol/PIP2 membrane condition shows a similar trend where the base compressibility modulus is observed to be ∼71 mN/m and increases to ∼80mN/m, which is a 12 % increase in the compressibility modulus. Remarkably, these changes in membrane compositions that were relatively less complex and devoid of essentially PI showed a reverse trend. Whereas in the case of DOPC/BSM/Chol 4:4:2 membrane condition, the base compressibility modulus was observed to be ∼ 84mN/m, which drastically decreased to approximately half of it, i.e., ∼ 41 mN/m on incubating it with Aβ-40. A∼ 50% decrease in compressibility modulus, which in simpler terms means a drastic increase in the monolayer compressibility. A similar trend in the DOPC/BSM/Chol 3:3:4 membrane condition is also observed where base compressibility modulus is ∼155 mN/m, and after incubation with Aβ-40 drops to ∼101 mN/m, which is a ∼ 30% decrease in the compressibility modulus.

### Aβ-40 fibril load generation on monolayers compressed at bilayer lateral pressure

After establishing the mixing behavior of Aβ-40 with free-standing monolayer and its effect on elasticity, we next wondered how would the Aβ-40 binding/fibrillation affect lipid monolayer membranes compressed to a surface pressure of 30-35 mN/m to mimic bilayer lateral pressure over the duration of fibrillation. The membrane monolayer was allowed to equilibrate for 15-20 minutes, after which the Aβ-40 was injected into the monolayer sub-phase (Fig. 7e). It was observed that immediately after injection of Aβ-40 within 1 hour, there was a drastic drop in the surface pressure of the myelin membrane from ∼30 mN/m to ∼2mN/m. This indicates that the peptide was inducing aggregation-based expulsion of the lipids into the sub-phase as it tried to populate the air/water interface already crowded by the lipid monolayer and, in the process, disrupting the membrane. Similarly, this was also observed in the case of DOPC/BSM/Chol/PIP2 but to a lesser degree, where the drop was about ∼16mN/m. In the case of both DOPC/BSM/Chol 4:4:2 and 3:3:4, there was a drop of ∼7mN/m and ∼2mN/m. Interestingly, the observed behavior of surface-pressure drop recapitulated our previous observations, wherein, Aβ-40 induced membrane disruption followed the similar order i.e, myelin membrane > DOPC/BSM/Chol/PIP2 > DOPC/BSM/Chol (4:4:2) > DOPC/BSM > DOPC/BSM/Chol (3:3:4). We reasoned that this could be because of the number of lipid packing defects available for Aβ-40 to embed itself at the interface. The much faster disruption of the membrane monolayers in comparison to the visible deformation in GUVs could be attributed to the lack of trans-bilayer interdigitation or leaflet coupling as well as the high sensitivity to the surface pressure changes in monolayer experiments.

Together, we show that the myelin model membrane can be deformed by the Aβ-40 mixture mimicking a high monomeric to oligomeric ratio. The deformation is driven by the interplay of the lipid packing defect densities mediated by early binding and the lipid-specific enhancement or retardation of the fibrillation of Aβ-40. The binding and fibrillation of Aβ-40 induce rigidification on short timescales followed by fluidization of the membrane over long time-scales, resulting in significant lipid extraction and tubulation. This increase in lipid extraction and tubulation with the Aß-40 fibrillation might hint at the possibility of biological demyelination during neurodegeneration (Fig. 8a-b).

**Fig. 8.**
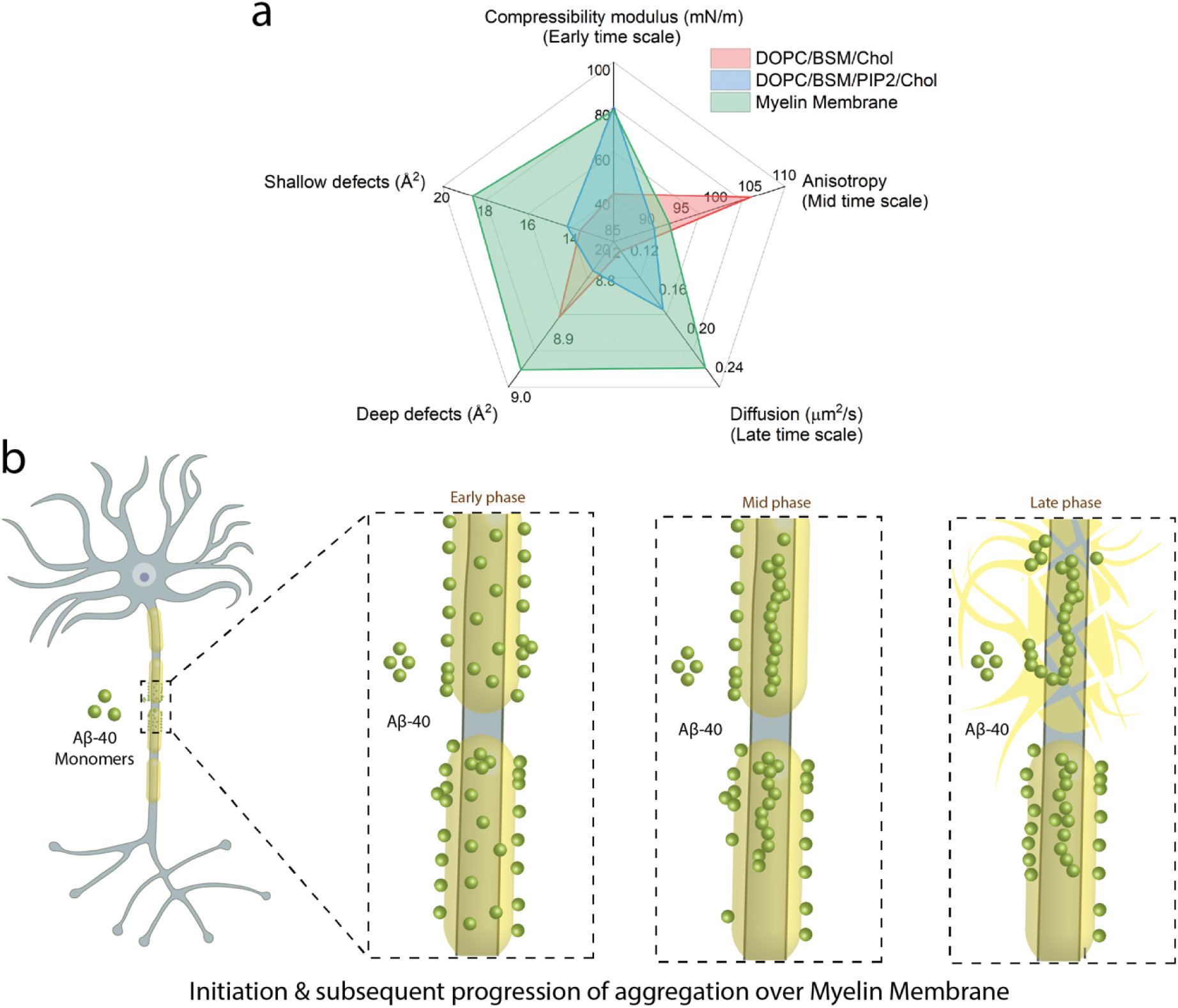
Correlation of changes in myelin membrane parameters induced by Aβ-40. **a)** Multivariate plot depicting the correlation between lipid defect densities, compressibility modulus, anisotropy and diffusion changes induced by Aβ-40 during its fibrillation. **b)** A schematic of the proposed model of progressive neuronal myelin deformation mediated by Aβ-40 aggregation. In the early phase, Aβ-40 binds and fluidizes the myelin membrane that subsequently facilitates the aggregation leading to disruption of the membrane.

## DISCUSSION

Emerging clinical evidence strongly suggests that myelin alteration is an important pathological feature in the pathogenesis of AD that may be one of the earliest characteristics of the disease progression (Dean *et al*., 2017). Despite the evidence that the soluble forms of Aβ are elevated in the white matter independent of the amyloid burden in the cortical plaque, whether Aβ aggregation can alter the myelin membrane remains elusive(Collins-Praino *et al*., 2014). In this study, we first reconstitute the binding of seeded Aβ-40 on the myelin lipid membrane, mimic and visualize snapshots of the changes at membrane interface over a period of 24 hours. The presence of a monomer/ oligomer mixture of Aβ-40 is known to accelerate the fibrillation through a physiologically more favored secondary nucleation mechanism, which is considered a major driving force during the progression of protein aggregation(Cohen *et al*., 2013). We see that the binding of Aβ-40 to the myelin membrane, although homogenous, shows an interesting oscillating pattern of increasing and decreasing binding intensity at early (1-4hour), mid (4-12 hour), and late phase (12-24 hour) (Fig. 1). The differences in binding might arise due to structural changes in the soluble forms of the Aβ aggregates within the heterogeneous population that exist in a highly dynamic metastable state. Such an ensemble of Aβ-40 forms is not only physiologically relevant but also results in different modes of interactions with the membranes(De *et al*, 2019). Extensive tubulation of the myelin membrane is observed in the late phase involving fibril-mediated phase separation within the membrane as a result of lipid sequestration by the growing fibril (Fig. 1). Aβ-40, particularly at the mid-late phase (4-12 hour), triggered the clustering of vesicles, likely entangled by the growing fibrils as also evident from the dynamic light scattering measurements showing the increased hydrodynamic radii due to the fusion/clustering of vesicles (Fig. 1, Fig. S12). Similar clustering of vesicles was reported earlier, induced by β_2_M fibrils, wherein ends of the fibrils were found to distort the membrane and extraction of lipids(Milanesi *et al*, 2012). Aggregation kinetics of Aβ-40 revealed that while myelin membrane enhances sustained aggregation of Aβ-40, however, except PIP2 other myelin lipid components such as DOPC, DOPG, PI, DOPS decrease the overall aggregation rate (Fig. 2d). There are contrasting observations previously reported on the effect of DOPC on Aβ fibrillation. While one study observed DOPC mediated retardation of Aβ-40 and Aβ-42 fibrillation(Hellstrand *et al*., 2010), another study reported enhancement of Aβ-42 fibrillation(Lindberg *et al*, 2017). The later study observed enhancement of DOPC driven Aβ-42 fibrillation was driven by augmentation of the secondary nucleation that facilitates fibril fragmentation. It was also observed that Aβ-42 monomers do not directly interact with DOPC membrane till fibrils are formed, something in line with our observation of delayed binding of Aβ-40 to DOPC membranes (Fig. 4a). While we could see significant binding of Aβ-40 to DOPG membrane however, it did not seem to have much effect on the aggregation kinetics (Fig 2a, 4d-e). This is in line with previous observations wherein POPC/POPG membrane had no effect on Aβ-40 fibrillation although DPPG induced templated crystalline ordering that triggers fibrillation(Chi *et al*., 2008). Interestingly, although we did not observe any significant binding of Aβ-40 to DOPS membrane in our study, however, Aβ-42 is shown to significantly interact with DOPS membranes in the presence of Ca^2+^ that help bridge Aβ-42 nucleate on the membrane(Yi *et al*, 2015). Likewise, the aggregation kinetics of another peptide amyloidogenic peptide IAPP was also found to enhance in the presence of higher mole % of DOPS(Zhang *et al*, 2017). Our observations on the influence of DOPE and PI membranes on Aβ-40 is in line with previous observations that suggested PE membranes hamper fibril formation in IAPP and that PI can help fibril elongation but not nucleation(McLaurin *et al*, 1998; Sciacca *et al*, 2012a). Such divergent modulation of the Aβ-40 binding and fibrillation by constituent lipids dissects the importance of lipid specificity in regulating the rate of Aβ-40 aggregation. The observation is in line with previously reported lipid-induced depolymerization of Aβ fibrils generating “reverse oligomers” that are also cytotoxic(Butterfield & Lashuel, 2010; Engel *et al*, 2008; Martins *et al*, 2008).

A key finding from our results is that the degree of binding by Aβ-40 depends on the interplay of lipid packing defects and electrostatics that eventually drive membrane deformation. Aβ-40 binds early when the membrane contains either a mixture of lipids with shapes that differ geometrically or possess a negatively charged surface (Fig. 2). The general trend for the average binding intensity of Aβ-40 during the early, mid, and late phase is Myelin membrane > PC/BSM/Chol/PIP2 > PC/BSM/Chol that reflect the decreasing density of the surface voids or the lipid packing defects at the membrane interface (Fig. 1, 3). In addition to the lipid packing defects other parameters such as net negative charge and the presence of cholesterol might also facilitate the observed binding(Yang *et al*., 2021; Zhang *et al*., 2017). DOPC membrane, although it contains conical lipids yet the lack of electrostatics results in no to weak binding of Aβ-40 (Fig. 4a). The geometrical arrangement of lipids with cylindrical and non-cylindrical shapes in myelin membrane and their entropically driven propensity to readily mix should result in the highest lipid packing defect density compared to membranes with less heterogeneous lipid shapes(Bigay & Antonny, 2012; Vanni *et al*, 2014). Further, the process of membrane deformation progresses through the fluidization leading to the disruption of the membrane as inferred from the anisotropy changes of the potentiometric dye (Fig. 4). The degree of observed fluidization depends on lipid packing density and complements the previous observation that reported Aβ-40 mediated fluidization of the neuronal membrane in a cholesterol-dependent manner (Ji *et al*., 2002; Yu & Zheng, 2012). Comparison of the state of binding/growth of Aβ-40 and its effect on lipid dynamics in mid to late phases (12h and 24h) showed that Aβ-40 is highly mobile on myelin membrane reflected in fluorescence recovery upon fluorescence photobleaching (Fig. 6). The mobility of Aβ-40 was also found to follow the same trend as the average binding, i.e., myelin membrane > PC/BSM/Chol/PIP2 > PC/BSM/Chol, suggesting that Aβ-40 has a highly dynamic interaction on membranes with higher densities of lipid packing defects. However, the strongest increase in the lipid mobility was observed in the case of the myelin membrane, followed by PC/BSM/Chol/PIP2 and least changed in the case of PC/BSM/Chol membrane (Fig. 6). This suggests that the growth of Aβ-40 on myelin membrane interface, although highly dynamic, however, the interaction with the underlying lipids is strong enough to reduce the average diffusion that increases over time due to membrane deformation (Fig. 6c, f; Table S1.). Aggregation of Aβ-40 on the membrane surface might result in the creation of diffusion barriers that would reduce lipid diffusion(Adrian *et al*, 2014; De Franceschi *et al*, 2019; Shrivastava *et al*., 2017). Such aggregation mediated reduction in lipid diffusion has also been observed for other amyloid proteins(Iyer *et al*, 2016).

The monolayer experiments helped us dissect the earliest timescales of molecular interaction of Aβ-40 with the membrane (i.e., within a time scale of 30mins to 1hour). The surface pressure-area isotherms of the myelin membrane show a significant increase in the area per lipid molecule around 30mN/m followed by a decrease in the collapse surface pressure hinting at Aβ-40 induced reduction in the compressibility (i.e., C_s_) of the membrane (Fig. 7). A similar observation was seen in the case of PC/SM/Chol/PIP2 membrane. On the contrary, Aβ-40 was found to induce a rise in collapse pressure in the case of PC/SM/Chol membranes (4:4:2 and 3:3:4 ratio), suggesting an overall increase in the compressibility (Fig. 7).

Furthermore, the Aβ-40 was found to significantly deform the monolayers of myelin and PC/SM/Chol/PIP2 membrane equilibrated at 30mN/m, as evident from the steep drop in the surface pressure within the early time regime (Fig. 7). The deformative effect seemed to be much less pronounced for PC/CM/Chol (4:4:2 and 5:5:0) while least in the case of the membrane with the highest amount of cholesterol (i.e., PC/SM/Chol (3:3:4). Our data suggest that the earliest binding of seeded Aβ-40 on the membranes strongly depends on the availability of the surface lipid packing defects. The striking reduction in the surface pressure can be attributed to the generation of load by the growing fibrils(Xue *et al*, 2009) or the extraction of lipids by the monomeric/oligomeric Aβ-40 and the lack of interleaflet coupling in the membrane monolayer(Harayama & Riezman, 2018). The above early time scale observations combined with the late time scale microscopic observations are in line with a recent report suggesting insertion of oligomer and protofibrils on liposomes observed by 3D Cryo-electron tomography(Tian *et al*, 2021). We think that although the monomeric or low oligomeric Aβ-40 may insert into the membrane depending on the depth of the lipid packing defect, however, the fibrillation takes place at the membrane interface, as evident from the lack of increase in surface pressure corresponding to surface insertion (Fig. 7) which is in-line with the purview that the deformation of a membrane in response to a force can be described on the basis of membrane compression, area expansion, and bending moduli(Tyler, 2012).

Taken together, the work captures both the molecular insights of both early and late events of the Aβ-40 mediated myelin membrane deformation. The findings demonstrate how lipid packing defects can be exploited by Aβ-40, to manipulate the compressibility modulus, anisotropy and diffusion of the myelin membrane to drive the aggregation mediated deformation (Fig. 8a). This may also be critical for the elusive mutual interference of the early pore formation and late fibril mediated lipid extraction that contribute to the two-step mechanism(Ding *et al*, 2012; Sciacca *et al*., 2012b; Sciacca *et al*., 2018; Wong *et al*, 2009). Interestingly, Alpha-Synuclein has been observed to sense lipid packing defects and induce lateral expansion of the lipids (Ouberai *et al*, 2013). The modulation of lipid packing and protein recruitment by changing the lipid composition has been shown important for several peripheral proteins (Vanni *et al*., 2014). The proposed biophysical mechanism of myelin deformation by Aβ-40 in vitro should draw attention towards the unexplored fundamental role of myelin in the onset and progression of neurodegeneration.

## Materials and Methods

1,2-Dioleoyl-sn-glycero-3-phosphocholine (DOPC), 1,2-dioleoyl-sn-glycero-3-phosphoethanolamine (DOPE), L-α-phosphatidylinositol (liver PI), 1,2-dipalmitoyl-sn-glycero-3-phosphocholine (DPPC), 1,2-dioleoyl-sn-glycero-3-phospho-Lserine (DOPS) 1,2-dioleoyl-sn-glycero-3-phospho-(1′-rac-glycerol) (DOPG), L-α-phosphatidylinositol-4,5-bisphosphate (PIP2), sphingomyelin (Brain, Porcine) (BSM), L-α-phosphatidylinositol-4,5-bisphosphate (Brain, Porcine), 1,2-dioleoyl-sn-glycero-3-phosphoethanolamine-N-(lissamine rhodamine B sulfonyl) (Rhod PE) and cholesterol were purchased from (Avanti Polar Lipids, Alabaster, Alabama, U.S.A.) Composition of myelin membrane - DOPC/BSM/DOPE/PI/DOPS/Cholesterol (4:3:1:1:0.4:0.6). Beta - Amyloid (1 - 40) (DAEFRHDSGYEVHHQKLVFFAEDV-GSNKGAIIGLMVGGVV), HiLyte™ Fluor 488 – labelled Beta - Amyloid (1- 40) (HiLyte™ Fluor 488-DAEFRHDSGYEVHHQKLVFFAEDVGSNKGAIIGLMVGGVV), Human, Anaspec (Fremont, CA, USA), Di-8-ANEPPS from Invitrogen™ Thermo Fisher Scientific was used for the fluorimetry experiments.

### Peptide reconstitution

The commercially available amyloid Beta 40 (Aβ 40) was purchased from AnaSpec, inc. which was stored at -20°C. At the time of preparation, the stored peptide was made to equilibrate at room temperature. The peptide powder was then dissolved in 40 μL of 1% NH_4_OH diluting it with milliQ water up to 1mL, bringing the concentration of the peptide at 1mg/mL. Further, 10μL aliquots of this preparation were flash freezed and lyophilized, which was then stored at -20°C. The peptide was then dissolved in the desired buffer for further experiments. The reconstituted peptide was incubated for two hours to allow aggregation prior to quantifying different populations of soluble forms of Aβ-40 by fluorescence correlation spectroscopy.

### Thioflavin T assay for the measurement of fibrillation Kinetics of Amyloid Beta 40

The freeze-dried peptide was then reconstituted in phosphate-buffered saline (PBS). The Final concentration of the peptide for ThT experiments was kept at 1μM. The Thioflavin T concentration used for the experiments was 20μM. The lipid specificity of Aβ 40 was screened by incubating the peptide with giant unilamellar vesicles (GUVs) which acted like lipid templates for amyloid aggregation. The fluorescence intensity was followed against time to monitor the Aβ 40 fibrillation kinetics using BioTek Synergy H1 fluorescence plate reader at an excitation wavelength of 440 nm and an emission wavelength of 490 nm. Readings in triplicate were recorded every 30 min for 6 hrs. To minimize evaporation an Opti-seal was applied over the microplate. The data was then normalised by the lipid controls for each condition and plotted using Origin pro. The initial growth rate was calculated by fitting the initial log phase of the aggregation kinetics to the equation y = A + B*exp(- kx).

### Preparation of fluorescently labelled large unilamellar vesicles

Each LUV sample contains 120 nmol of the respective lipid which is doped with 1 mol% of di-8-ANEPPS. This lipid solution was then dried under a gentle nitrogen gas stream, subsequently it was vacuum dried for an hour to remove the residual solvent from the lipid film. These lipid films were then rehydrated in 250 μL of PBS of pH 7.4 and then were incubated for about 15 min in a water bath, making sure the temperature remained above the transition temperature of the lipid. The heated samples were then vortexed for 4-5mins. For the preparation of LUVs, the MLV suspension was then sonicated for 5 min at 0.9 pulse rate and 100% amplitude. The size of the LUVs was confirmed using DLS with the average diameter of the LUVs being ∼ 200 nm. Lipids used for background correction were devoid of dye.

### Preparation of fluorescently labelled giant unilamellar vesicles (GUVs)

The gel-assisted described by Weinberger et al. was followed for the preparation of GUVs. Briefly, a 5%(w/w) solution of Poly-vinyl alcohol (PVA) was prepared in deionised water, 300 μL of this solution was evenly spread on a plastic petri dish and dried at 50°C for 30 min in an oven. From a solution of lipids in chloroform at 1 mg mL^−1^ concentration doped with 1 mol% rhodamine-PE, 20 μL of this solution was spread on the PVA coated petri dish. These Petri dishes were then placed under vacuum for 45 min to sufficiently dry the lipid film. To prevent dewetting, the petri dishes were cleaned with UV for 15 min. This lipid film was then left to swell in phosphate buffered saline (PBS) at a pH of 7.4 for 45 min. The hydrated vesicles were gently dislodged and transferred to a microcentrifuge tube using a pipette(Tiwari *et al*, 2018).

### Confocal fluorescence microscopy

A custom-made chamber was used for incubating the GUVs and Aβ 40 doped with 10% Hylite-488 Aβ 40. The cover slip was wipe cleaned with 70% ethanol and was then air-dried. Then, 90 μL of the GUVs from the microcentrifuge tube was added to the chamber along with 10 μL of Aβ 40 in equi-osmolar PBS buffer with the effective concentration of the peptide being 2.5μM. This chamber was then sealed with an opti-seal so as to minimize evaporation during prolonged incubation of the sample. Imaging was performed on a Leica TCS-SP8 confocal instrument using appropriate lasers for rhodamine-PE (DPSS-561) and Hylite-488 Aβ 40 (argon-488). An identical laser power and gain settings were used during the course of all the experiments. The image processing was done using ImageJ.

### Quantification of Membrane Packing Defects

The biological membranes are composed of lipids of different shapes such as cylindrical, conical etc. and when such lipids get together to frame a bilayer structure with their polar heads exposed to the aqueous environment, several voids get created which leads to the packing defects in membrane. To study these defects, membranes of four different compositions were set up and simulated using coarse-grained (CG) molecular dynamics. The CG simulations were performed using the MARTINI version 2.2(Marrink *et al*, 2007) force field in GROMACS version 5.1(Abraham *et al*, 2015). The four membrane systems with the varying compositions are: 1) pure DOPC(dioleoyl-phosphatidylcholine), 2) DOPC:BSM(brain sphingomyelin):CHOL at a molar ratio of 4:4:2, 3) DOPC:BSM:CHOL:PIP2(Phosphatidylinositol bis-phosphate) at a molar ratio of 2:4:3:1, 4) myelin membrane DOPC:BSM:CHOL:PI(Phosphatidylinositol):DOPE:DOPS at a molar ratio of 4:3:1:1:0.4:0.6. The python script, insane.py, was used to generate each of the above mentioned membrane systems within a simulation box of 12 x 12 x 10 nm^3^ solvated with explicit MARTINI water and with appropriate numbers of Na+ and Cl^-^ counter ions to make the charge of each of the system neutral(Van Hilten *et al*, 2020). After running a steepest descent minimization, a 200 ns NVT equilibration was executed with the temperature coupling by velocity rescaling thermostat to a reference temperature of 300K. The NVT equilibration is followed by a 200 ns NPT equilibration with the pressure coupling by Parinello-Rahman barostat to a reference pressure of 1 bar. After the equilibration, a 10 μs production run for each of the bilayer system was performed out of which the last 1 μs was used for analysis.

We used PackMem(Gautier *et al*., 2018) to quantify the lipid packing defects. PackMem follows the Cartesian grid system for mapping of the membrane surface where the grid dimension is set to 1 Å x 1 Å. This tool computes the defects by characterizing it into deep and shallow defects. The deep defects represent the voids created due to the presence of aliphatic atoms deeper than d Å (where the value of d is set to 1Å) below the central atom of glycerol whereas shallow defects represent the accessible aliphatic atoms that are less than d Å below the central atom of glycerol and all types represent the combination of both the deep as well as shallow defects ^[2]^. For the execution of PackMem, the MARTINI trajectory files obtained from the production run were taken as input to generate the pdb frames saved every 200 ps. The script “ScriptPackMem.sh” then takes each pdb file as input to calculate deep and shallow defects. The distribution of packing defects area follows a mono-exponential decay,

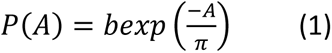

where *P(A)* denotes the probability of finding a defect with area *A*, *b* is the pre-exponential factor and *π* is the packing defect constant(Gautier *et al*., 2018). Finally, an R script provided with the package computes the mean packing defect constants. The barplot for the packing defect constants for each type of packing defects is prepared using GNUPLOT version 5.2(Williams *et al*, 1986).

### Fluorescence recovery after photobleaching (FRAP)

For the FRAP measurements on the GUVs, the GUVs were doped with 1 mol% of rhodamine PE and were incubated with 2.5 μM of Aβ 40 doped with 10% Hylite-488 Aβ 40. First, pre-bleach images at an attenuated laser intensity were acquired. Photobleaching was performed using DPSS-561 (to photo-bleach the lipid rhodamine-PE) and argon-488 (to photo-bleach bound the bound Hylite-488 labelled Aβ 40) at 100% laser power for 30 s, achieving a partial bleach through a circular region of interest (ROI) of a nominal radius r = 2.2–2.4 μm. The laser was then switched back to the attenuated intensity, and the recovery curve along with the images was recorded for several seconds. The photobleaching was executed at the equatorial plane of the GUV being visualised. The FRAP curves for each condition was repeated five times and then normalized. The diffusion coefficient was calculated using the Soumpasis equation for 2D-diffusion

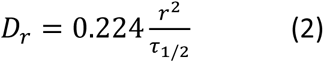

Where, 0.244 is the numerically determined value, r (2.2 μm) stands for the radius of the laser beam focused on the region of interest, and τ_1/2_ is the time required for half the recovery. The time for half the recovery was determined by plotting the normalized recovery curve.

### Fluorescence spectrophotometric assay

The Fluorescence spectroscopy experiments were performed on LUVs (preparation described earlier). The LUVs prepared were then incubated with Aβ 40 at an effective concentration of 1μM and making up the total volume of the LUV including the Aβ 40 to 300 μL. This mix was then incubated in dark. Readings were then taken at the 4^th^ hour of incubation of aggregation kinetics of Aβ 40. The fluorescence anisotropy was measured for 24s, with an integration time of 1s. 460nm and 560nm were the excitation and emission wavelength, respectively. The anisotropy (r) was automatically calculated by the instrument using the equation;

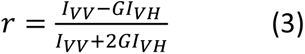

where,

I_VV_ and I_VH_ are the measured fluorescence intensities with the excitation polarizer oriented vertically and the emission polarizer oriented vertically and horizontally, respectively. G (=I_HV_/ I_HH_) is the grating correction factor that corrects for wavelength-dependent distortion of the polarizer. All experiments were conducted with multiple sets of samples. The above experiments were performed on a Perkin Elmer LS-55 steady state Fluorimeter.

### Fluorescence Correlation Spectroscopy (FCS) Experiments and Data Analysis

FCS experiments were carried out using a dual channel ISS Alba V system equipped with a 60X water-immersion objective (NA 1.2). Samples were excited with an argon laser at 488 nm. All protein data were normalized using the *τ_D_* value obtained with the free dye (Alexa488) which was measured under identical conditions. For a single-component system, diffusion time (*τ_D_*) of a fluorophore and the average number of particles (*N*) in the observation volume can be calculated by fitting the correlation function [*G(τ)*] to Eq.1 :

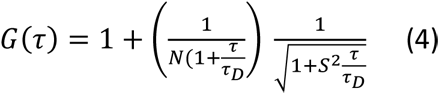

where, *S* is the structure parameter, which is the depth-to-diameter ratio. The characteristic diffusion coefficient (*D*) of the molecule can be calculated from *τ_D_* using Eq. 2:

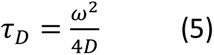

where, *ω* is the radius of the observation volume, which can be obtained by measuring the *τ_D_* of a fluorophore with known *D* value. The value of hydrodynamic radius (*r_H_*) of a labelled molecule can be calculated from *D* using the Stokes-Einstein equation [Eq. 6]:

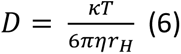

where, *k* is the Boltzmann constant, *T* is the temperature and *η* corresponds to the viscosity of the solution.(Chattopadhyay *et al*, 2002)

### Transmission Electron Microscopy

100μL of 200μM LUV solution was incubated with 1 μM Aβ 40 for different chosen time points. 10 μL of the incubated sample was then added on to a carbon coated copper grid and this was left for 2 minutes, it was later wicked off with a filter paper. The grid was then rinsed with deionized water and a 5 μL 4% uranyl acetate replacement (EMS) droplet was placed on to the grid. After minutes, this solution was wicked off and the grid was air dried. The imaging was performed on a JEOL (JEM 2100F) microscope with an operating voltage of 200 kV.

### Monolayer experiments

Langmuir monolayer films were spread on a Teflon molded trough (Apex Instruments, India) having inner working dimension of 305 mm X 105 mm. The rectangular trough was equipped with two movable Teflon barriers that provide a symmetrical compression. The system is equipped with an electronic balance having a sensitivity of ±0.5 mN/m to measure changes in surface pressure with the help of a suspended Wilhelmy plate (filter paper of dimensions 10 x 25 mm^2^). The entire system was set inside a transparent glove box. Before each experiment, the trough was cleaned with methanol, ethanol and ultrapure water so as to minimize impurities on the surface of water. PBS was used as the subphase which was maintained at a temperature of 25°C. Lipid solution of 1 mg/mL concentration was spread gently over the air/water interface until a surface pressure of 2-3 mN/m was reached and was left undisturbed for 15 min so as to relax the monolayer to 0 mN/m. 10 µL of Aβ 40 (23 µM) was injected into the subphase prior to monolayer compression. The subphase was gently stirred using a magnetic stirrer. Surface pressure(π)-Molecular area(A) Isotherms were recorded by compressing the monolayer at a constant speed of 3 mm/min. Isotherms were recorded until collapse pressure (π_c_) was reached. Isotherm data was used to process the compressibility modulus (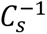)(Allende *et al*, 2004; Brockman, 1999; Brown & Brockman, 2007; Smaby *et al*, 1996)

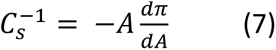

and bending force (Δ*F_b_*).

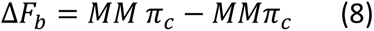

where, MMAβπ_c_ is the collapse pressure of model membrane with Aβ in subphase and MMπ_c_ being without Aβ. Negative values of (ΔF_b_) suggest bending of monolayer(Peetla *et al*, 2014).

For constant area measurements (Time v/s Pressure), lipid monolayer was first compressed to a surface pressure of 30 mN/m and left to relax for 10 minutes. Aβ 40 peptide was injected from underneath one of the barrier so as to avoid disturbing the monolayer and subsequently recorder for changes in surface pressure over time.

### Image processing

ImageJ’s oval profile plugin was used to extract intensity data from circumference of GUVs imaged under confocal microscope using along oval analysis option. Tubeness plugin was used to resolve the fibril like structures present in the fluorescence images of the GUVs. The plugin utilizes the eigenvalues of the Hessian matrix to calculate the measure of “tubeness” in case of 2D images, if the larger eigenvalue is negative, an absolute value is returned otherwise its returned as 0.

## ACKNOWLEDGEMENT

We gratefully acknowledge Sushant Pradhan, Chahat Kausar, Tapas Ghosh for technical assistance. AT acknowledge MHRD, India, for GATE-fellowship. We thank Dr Hirak Chakraborty for stimulating discussion. MS thanks the financial support provided in grant EMR/2017/004513 by the Department of Science and Technology (SERB-DST) and grant BT/PR21226/MED/122/41/2016 by the Department of Biotechnology (DBT), Government of India.

## AUTHOR CONTRIBUTION

MS and AT conceptualised the project. MS, AT designed the experiments and SJ helped with analysis of some experiments. AT performed most of the experiments. SP and MB performed the simulations. AS and KC performed the FCS analysis. AT made the figures. MS and AT wrote the manuscript with inputs from all the authors.

## Supplementary Information

**Fig. S1.**
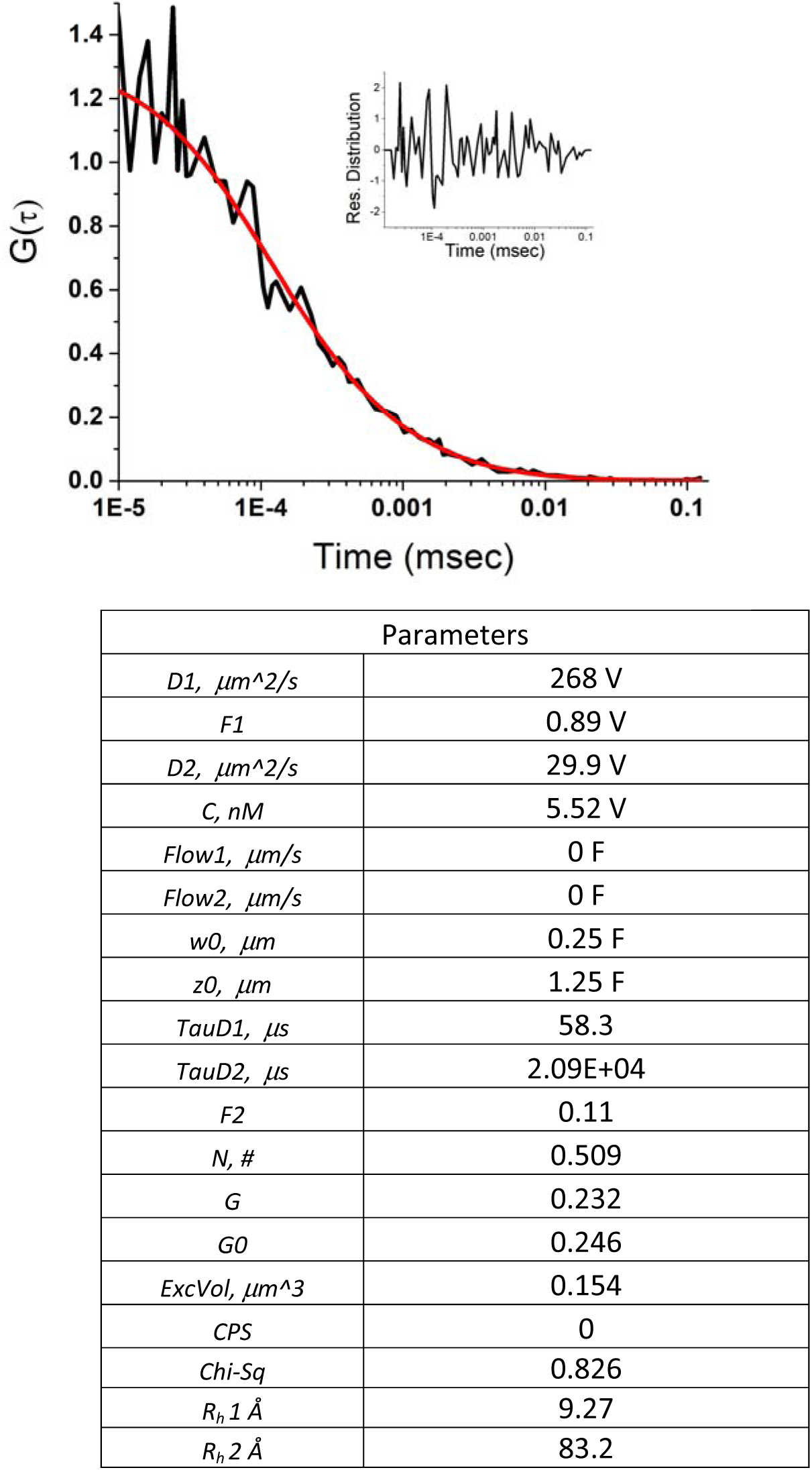
Fluorescence correlation spectroscopy (FCS) measurements to screen the population of Aβ-40 species present in the sample at a 0-hour time point.

**Fig. S2.**
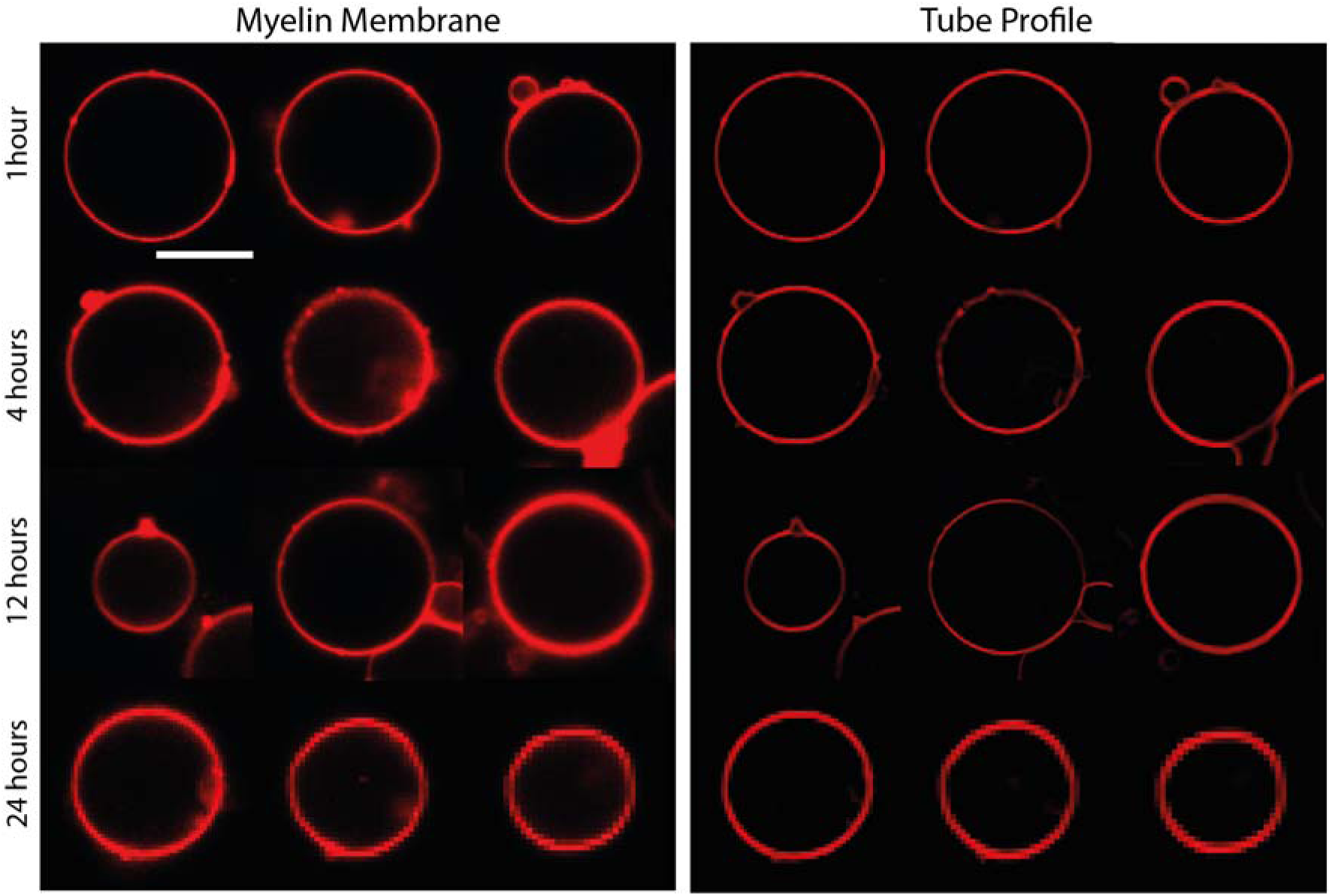
Representative images of GUVs of myelin membrane without Aβ-40 at respective time points. Scale bar is 10 μm.

**Fig. S3.**
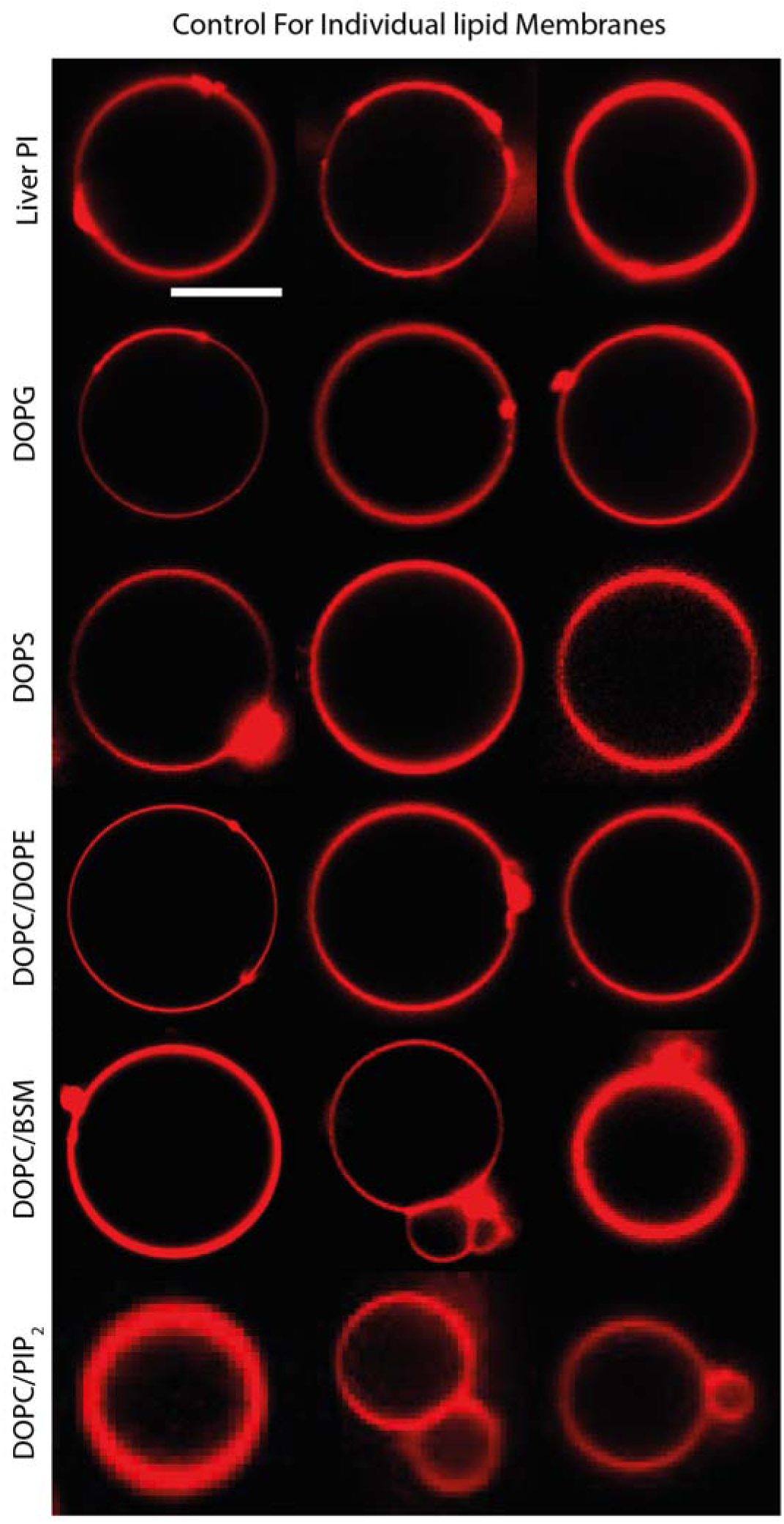
Representative images of GUVs of different membrane conditions at a 1-hour time point. Scale bar is 10 μm.

**Fig. S4.**
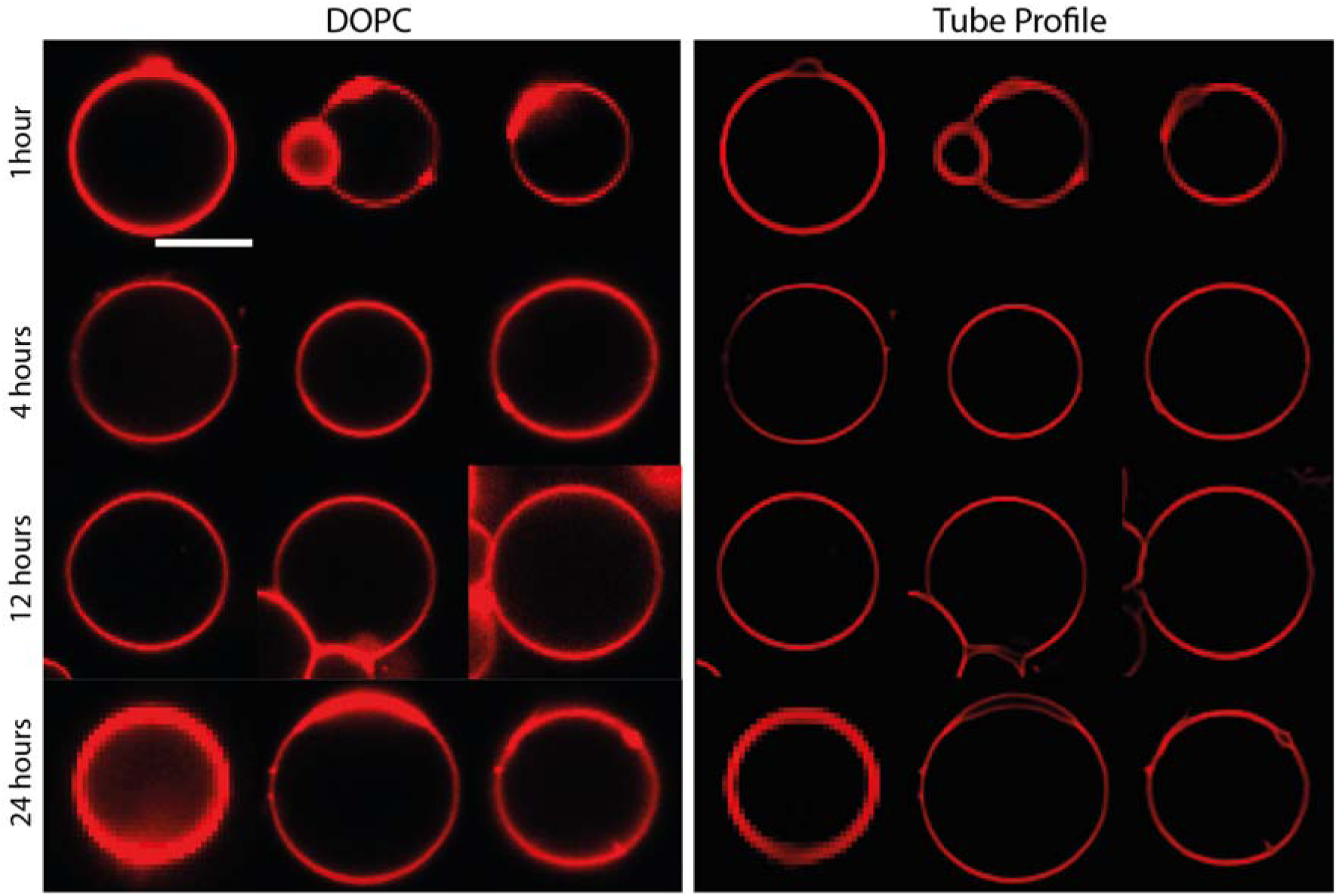
Representative images of GUVs of DOPC membrane without Aβ-40 at respective time points. Scale bar is 10 μm.

**Fig. S5.**
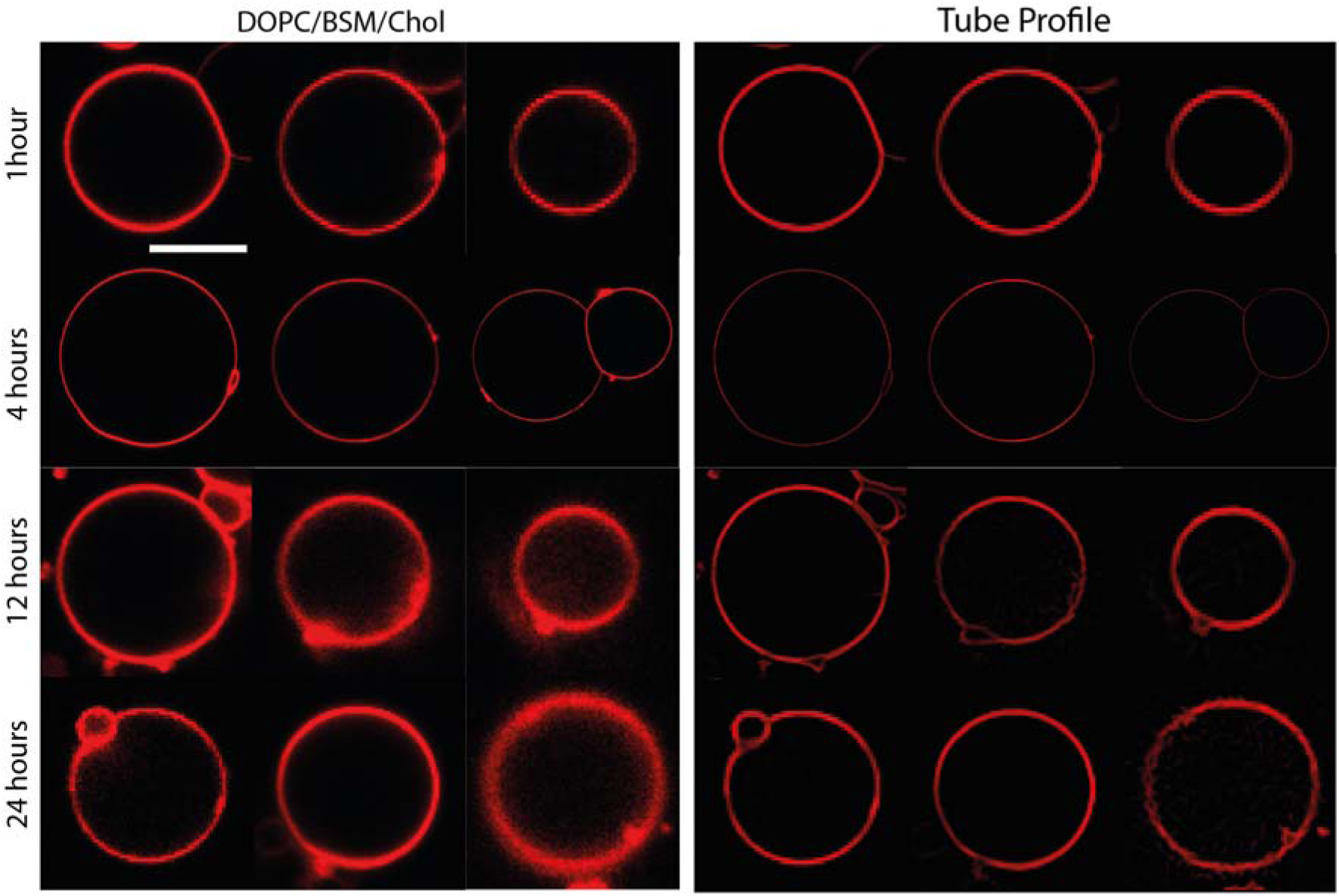
Representative images of GUVs of DOPC/BSM/Chol (4:4:2) membrane without Aβ-40 at respective time points. Scale bar is 10 μm.

**Fig. S6.**
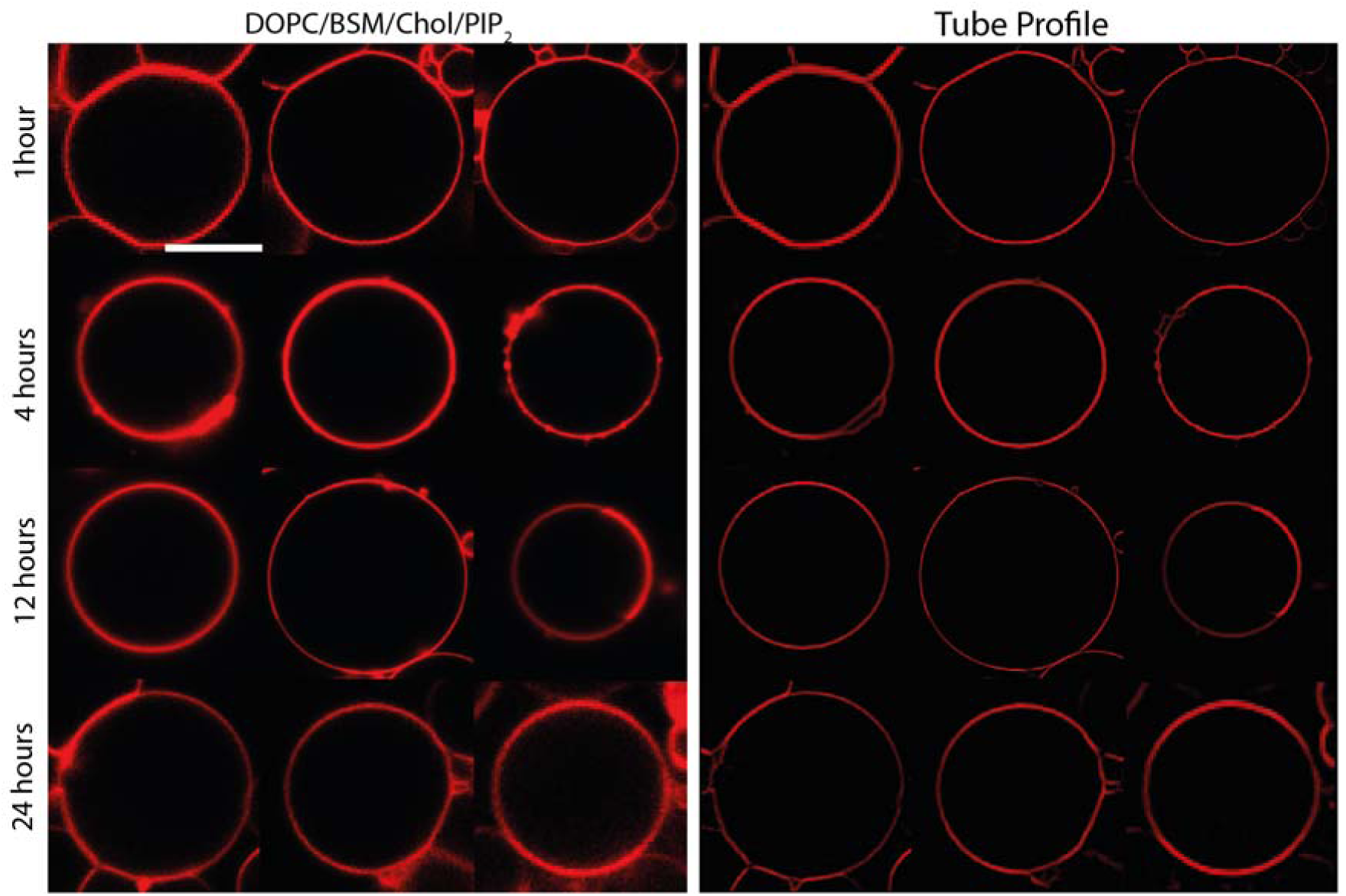
Representative images of GUVs of DOPC/BSM/Chol/PIP_2_ (2:4:3:1) membrane without Aβ-40 at respective time points. Scale bar is 10 μm.

**Fig. S7.**
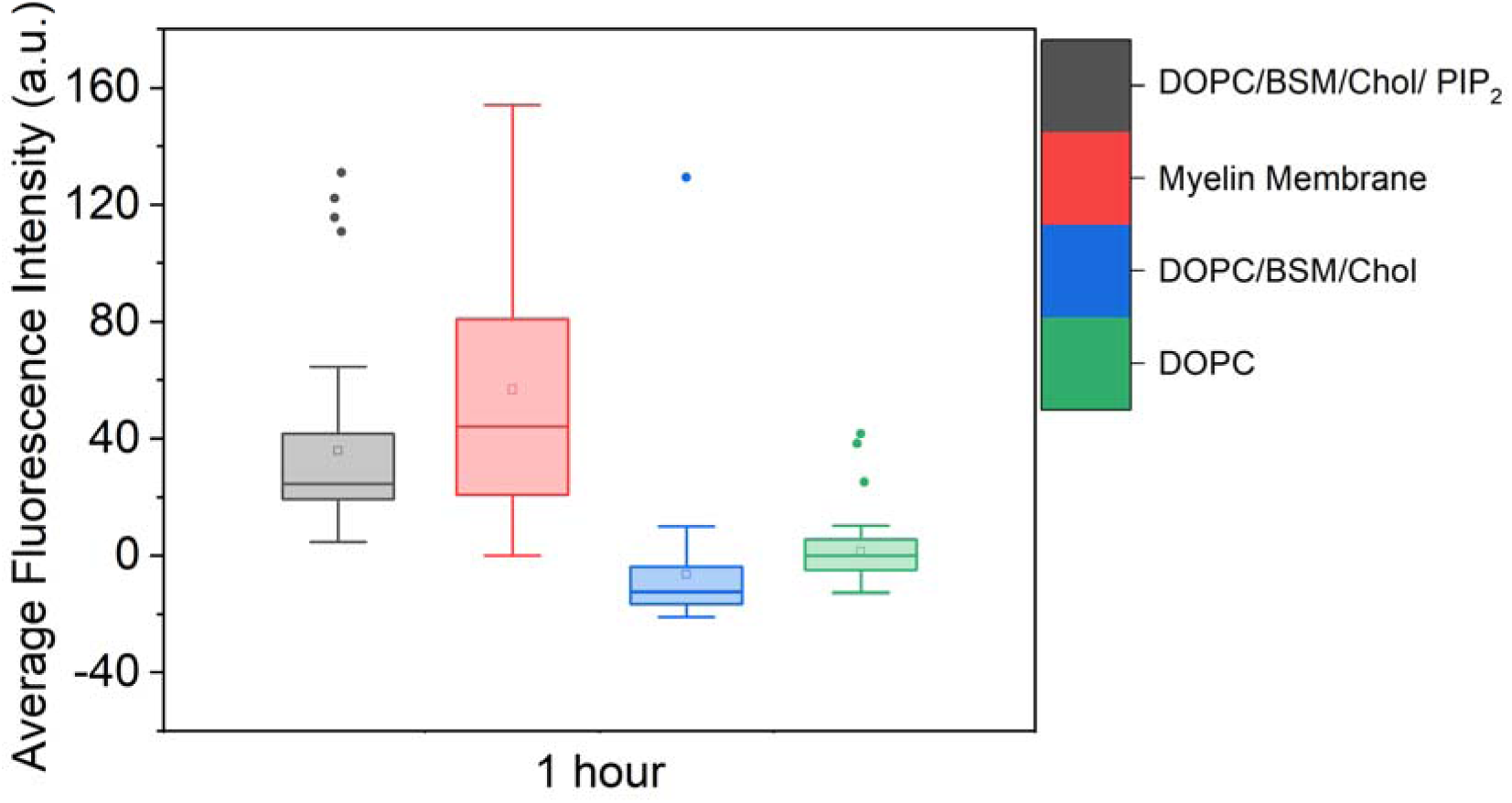
Box plot of the average binding intensity of Aβ-40 at the 1-hour time point for different membrane conditions.

**Fig. S8.**
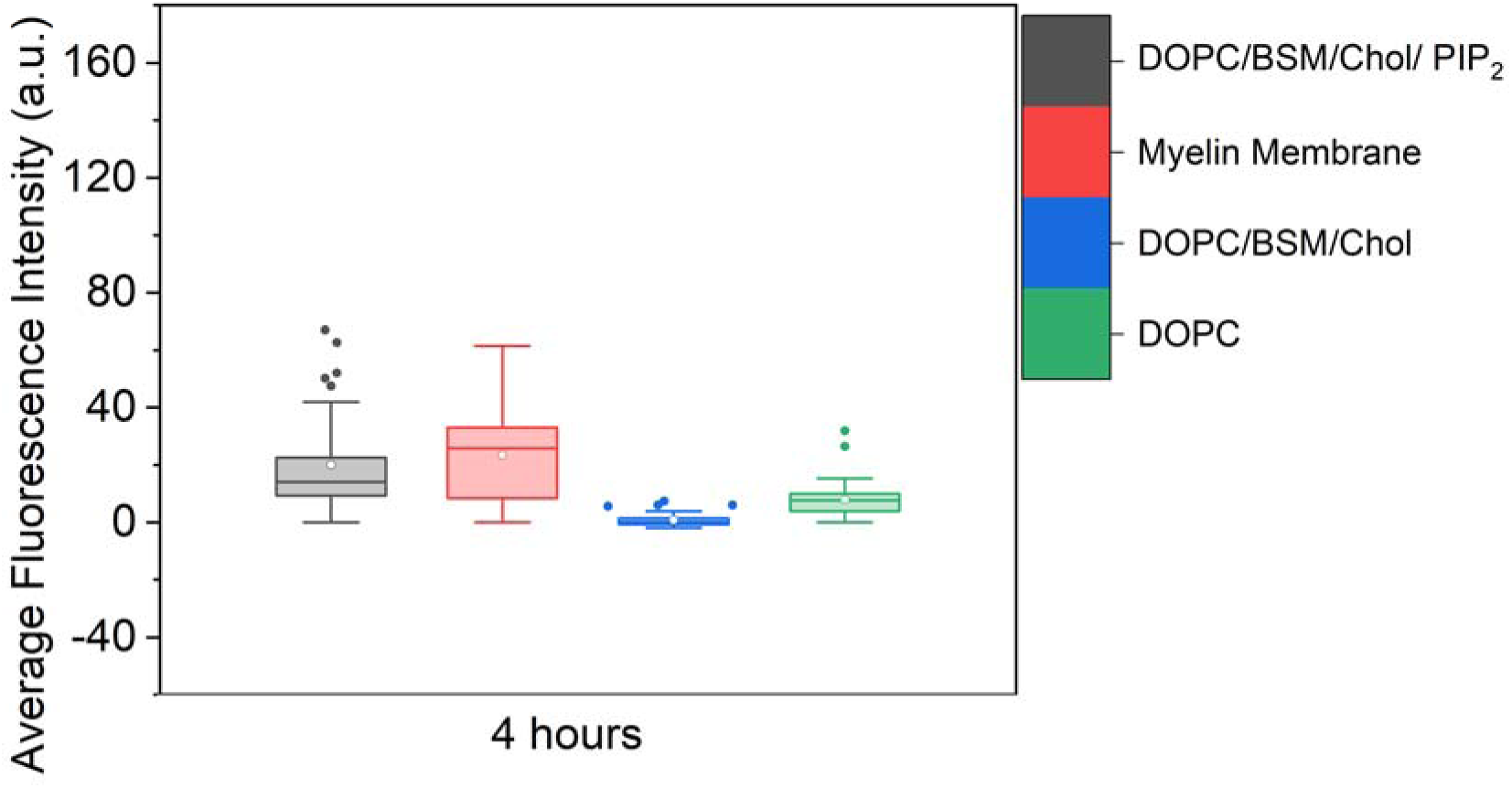
Box plot of the average binding intensity of Aβ-40 at the 4-hour time point for different membrane conditions.

**Fig. S9.**
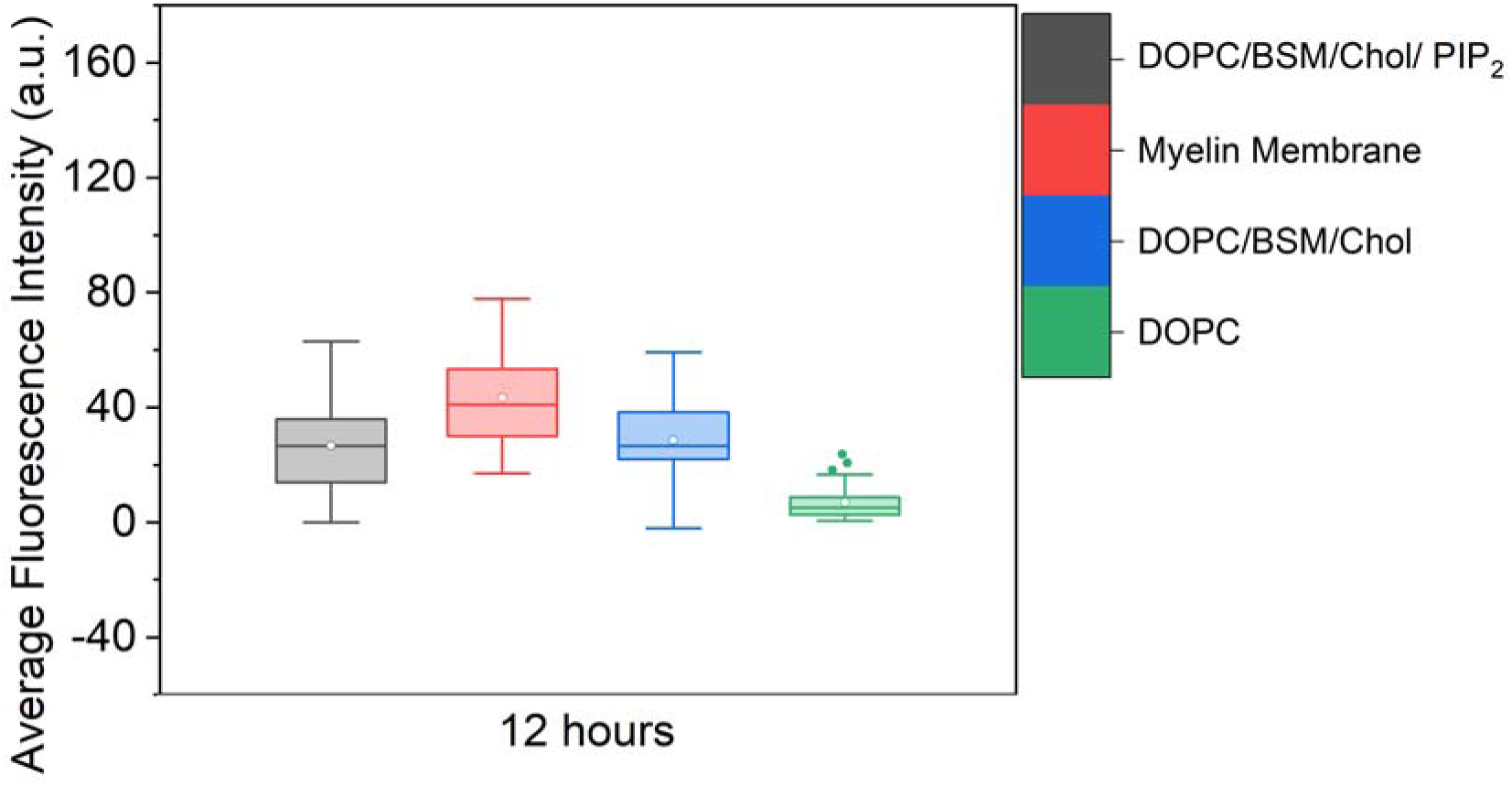
Box plot of the average binding intensity of Aβ-40 at the 12-hour time point for different membrane conditions.

**Fig. S10.**
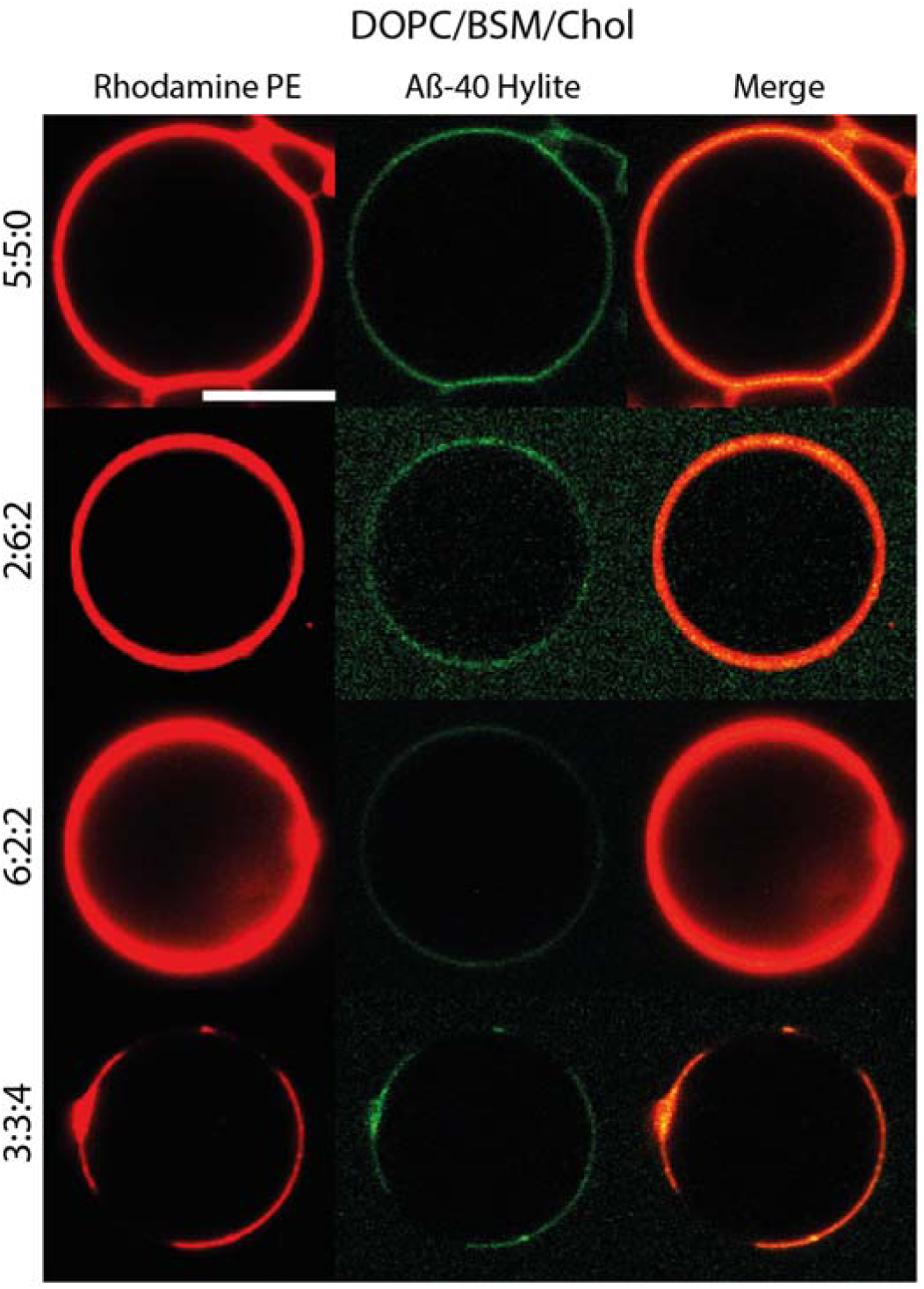
Interplay of ratio of lipid shape in a ternary membrane condition containing cholesterol on amyloid binding. Scale bar is 10 μm.

**Fig. S11.**
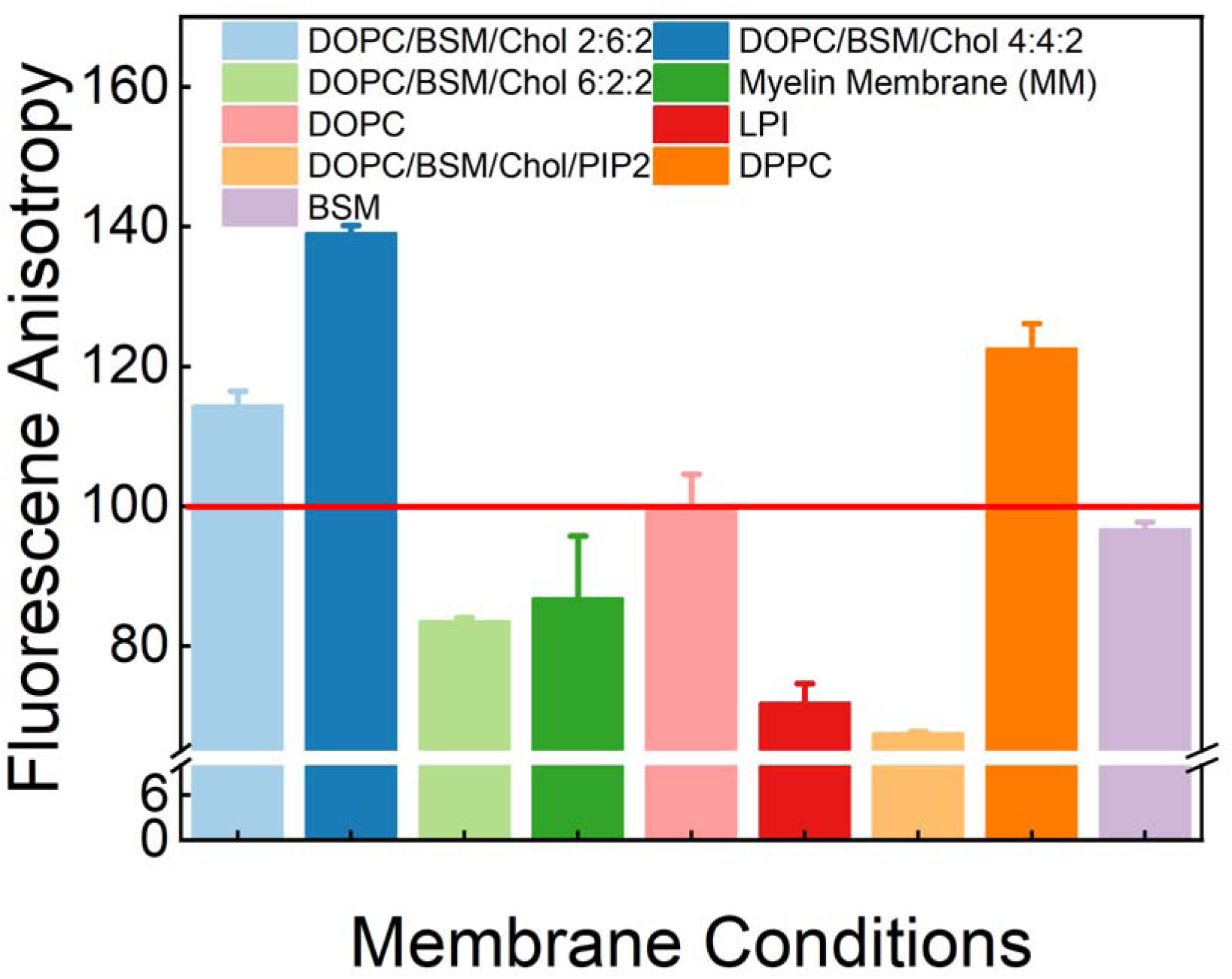
Steady-state anisotropy for different membrane conditions, controls without Aβ-40 for each membrane condition were normalized to 100 marked by the horizontal red line traversing the plot.

**Fig. S12.**
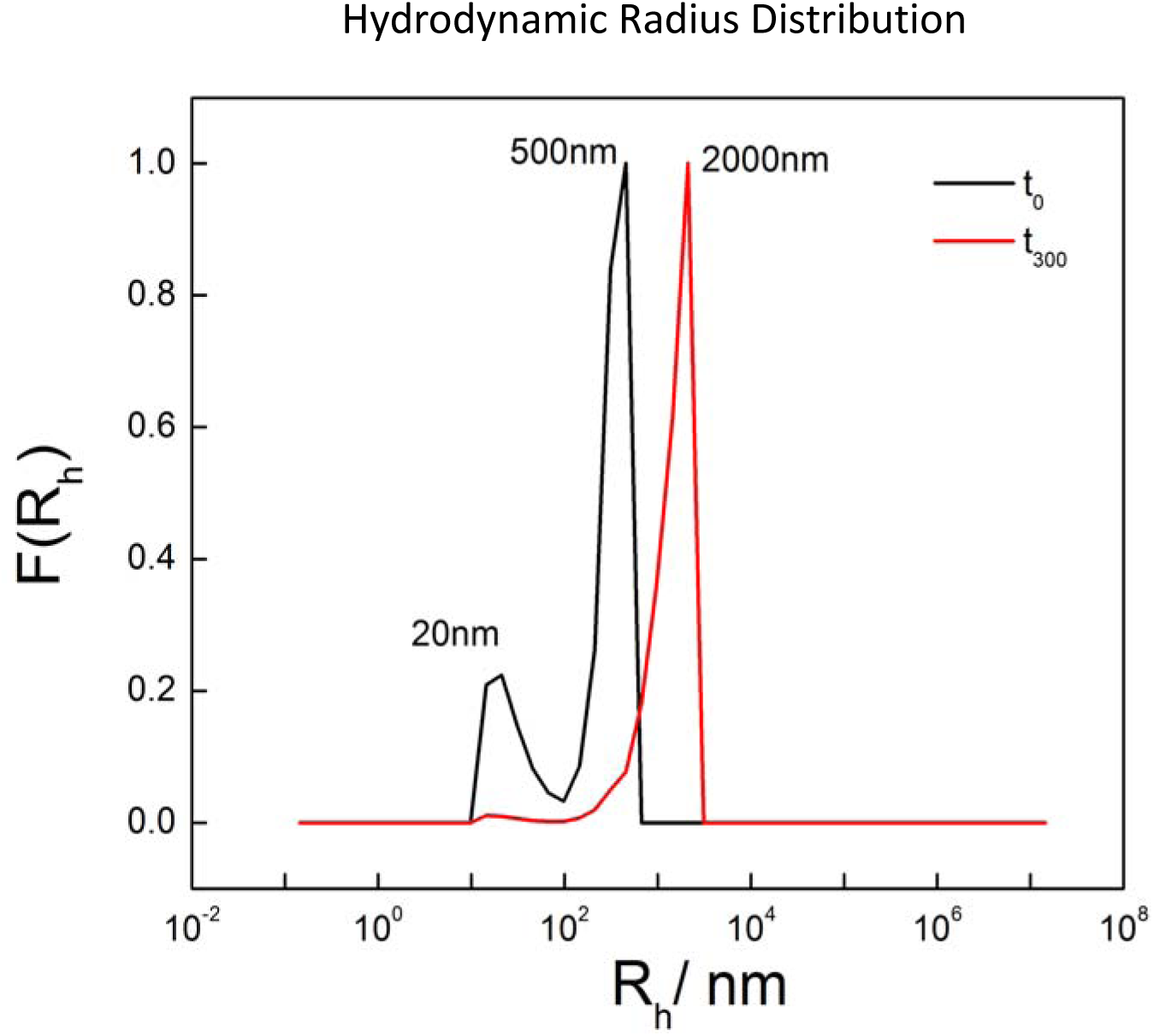
Dynamic light scattering measurement of DOPC LUVs incubated with Aβ-40 showed an increase in size at a 5-hour time point, indicating LUV-LUV fusion.

**Table S1.** FCS Table

| Time (hours) | D1 ($\mu\text{m}^2/\text{s}$) | D2 ($\mu\text{m}^2/\text{s}$) | TD1 ($\mu\text{s}$) | TD2 ($\mu\text{s}$) | F1 | F2 |
| --- | --- | --- | --- | --- | --- | --- |
| 0 | 268.75 | 29.9 | 58.3 | 20700 | 0.89 | 0.11 |
| 4 | 383.20 | 53.66 | 40.77 | 23143 | 0.67 | 0.33 |
| 8 | 330.19 | 55.14 | 47.32 | 24304 | 0.62 | 0.38 |

**Table S2.**
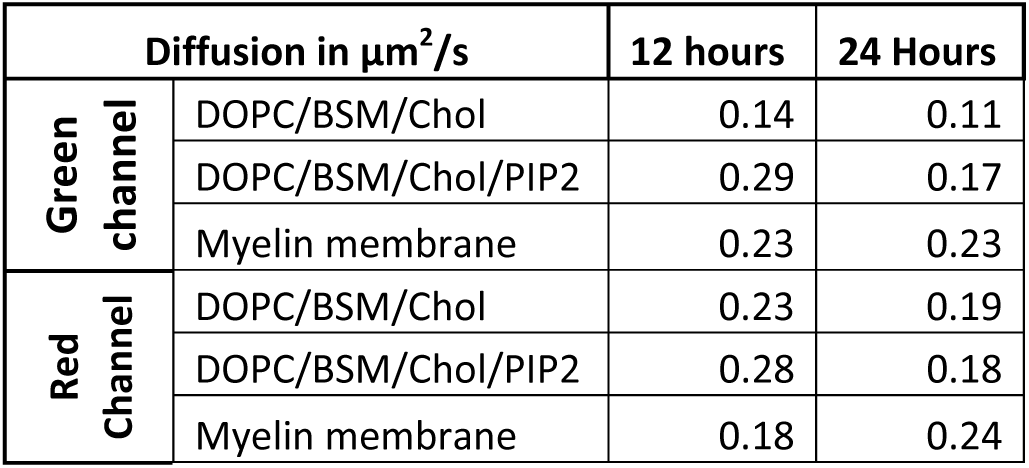
FRAP Diffusion Table

| Diffusion in $\mu\text{m}^2/\text{s}$ | | 12 hours | 24 Hours |
| --- | --- | --- | --- |
| Green<br>channel | DOPC/BSM/Chol | 0.14 | 0.11 |
|  | DOPC/BSM/Chol/PIP2 | 0.29 | 0.17 |
|  | Myelin membrane | 0.23 | 0.23 |
| Red<br>Channel | DOPC/BSM/Chol | 0.23 | 0.19 |
|  | DOPC/BSM/Chol/PIP2 | 0.28 | 0.18 |
|  | Myelin membrane | 0.18 | 0.24 |

